# Nitrogen cycling activities during decreased stratification in the coastal oxygen minimum zone off Namibia

**DOI:** 10.1101/2022.11.15.516596

**Authors:** Aurèle Vuillemin

## Abstract

Productive oxygen minimum zones are regions dominated by heterotrophic denitrification fueled by sinking organic matter. Microbial redox-sensitive transformations therein result in the loss and overall geochemical deficit in inorganic fixed nitrogen in the water column, thereby impacting global climate in terms of nutrient equilibrium and greenhouse gases.

Here, geochemical data are combined with metagenomes, metatranscriptomes and stable-isotope probing incubations from the water column and subseafloor of the Benguela Upwelling System. Taxonomic assemblages and relative expression of functional marker genes are used to explore metabolic activities by nitrifiers and denitrifiers under decreased stratification and increased lateral ventilation in Namibian coastal waters. Active planktonic nitrifiers were affiliated with *Candidatus* Nitrosopumilus and *Candidatus* Nitrosopelagicus among Archaea, and *Nitrospina, Nitrosomonas, Nitrosococcus*, and *Nitrospira* among Bacteria. Concurrent evidence from taxonomic and functional marker genes shows that populations of Nitrososphaeria and Nitrospinota were highly active under dysoxic conditions, coupling ammonia and nitrite oxidation with respiratory nitrite reduction, but minor metabolic activity towards mixotrophic use of simple nitrogen compounds. Although active reduction of nitric oxide to nitrous oxide by Nitrospirota, Gammaproteobacteria and Desulfobacterota was tractable in bottom waters, the produced nitrous oxide was apparently scavenged at the ocean surface by Bacteroidota. Planctomycetota involved in anaerobic ammonia oxidation were identified in dysoxic waters and their underlying sediments, but were not found to be metabolically active due to limited availability of nitrite. Consistent with water column geochemical profiles, metatranscriptomic data demonstrate that nitrifier denitrification prevails over canonical denitrification when Namibian coastal waters are ventilated during austral winter.

## Introduction

Oxygen minimum zones (OMZs) represent hotspots for oxygen-sensitive nitrogen microbial transformations (Codispoti and Christensen, 1985; Bange et al., 2000) and are traditionally viewed as productive areas dominated by heterotrophic denitrification fueled by algal organic matter (OM) sinking from the sunlit ocean surface down to the seafloor (Ulloa et al., 2012, 2013). In coastal waters such as the eastern tropical Pacific (Lam et al., 2009; Canfield et al., 2010), the Arabian Sea (Newell et al., 2011; Orsi et al., 2017), and the Benguela upwelling (Lam et al., 2009; Callbeck et al., 2021), oxygen drawdown in OMZ waters initiate a dynamic nitrogen cycle (Kuypers et al., 2005; Ulloa et al., 2012) in which nitrate serves as the main terminal electron acceptor for the oxidation of OM, is actively reduced to nitrite (Lam and Kuypers, 2011) and successively converted to nitrogen (N_2_) and nitrous oxide (N_2_O) gases through processes of heterotrophic denitrification (Tyrrell and Lucas, 2002) and autotrophic anaerobic ammonium oxidation (anammox) (Kuypers et al., 2005; Woebken et al., 2007). Dissimilatory nitrate reduction to ammonium (DNRA), or “ammonification” (Dong et al., 2009; Smith et al., 2015), is considered more nitrogen conservative (Jensen et al., 2011) as the resulting release of ammonium (NH_4_^+^) regenerates nitrate via microaerobic microbial respiration in the upper part of the OMZ (Fernandes et al., 2012; Behrendt et al., 2013) and surface oxygenated waters (Kalvelage et al., 2011, 2015; Füssel et al., 2012). Although all three microbial processes (i.e. denitrification, DNRA, anammox), compete for nitrate and nitrite (Dong et al., 2009; Kraft et al., 2011), the uptake of nitrate and its reduction to ammonia inside the cell and excretion via DNRA may achieve higher growth yields if pursued by ammonia oxidation and denitrification (Lam and Kuypers, 2011; Li et al., 2013). In general, anaerobic processes in the water column result in an overall geochemical deficit in inorganic fixed nitrogen and its loss from the oceans globally impacts the Earth climate system in terms of nutrient equilibrium (Küster-Heins et al., 2010; Lomnitz et al., 2016) and emissions of greenhouse gases (Codispoti et al., 2001; Doney et al., 2004; Kuypers et al., 2005; Bertagnolli and Stewart, 2018). OMZs are predicted to expand and intensify in response to global climate change and altered ocean circulation (Garçon et al., 2019), leading to increase in carbon sedimentation to the seafloor and benthic releases of greenhouse gases mediated by the resident microbial communities (Wright et al., 2012). It is therefore of utmost importance to disentangle metabolic interactions in biogeochemical cycling (Inthorn et al., 2006b; Löscher et al., 2016) within productive coastal OMZ waters and their underlying sediments.

Due to natural eutrophic conditions, the sunlit ocean surface of the Benguela Upwelling System (BUS) off Namibia is highly productive (Inthorn et al., 2006b; Verheye et al., 2016), leading to oxygen depletion below 60 m water depth (mwd) and a seasonal OMZ reaching down to the seafloor in austral summer (Lavik et al., 2009; Callbeck et al., 2021). Anoxic conditions extending down to the sediment-water interface (SWI) in austral summer trigger benthic releases of sulfide (H_2_S), methane (CH_4_) and NH_4_^+^ from the seafloor that seasonally intensifies OMZ conditions (Brüchert et al., 2009; Schunck et al., 2013; Neumann et al., 2016; Ohde and Dadou, 2018). Due to the quasi-absence of sedimentary Fe to precipitate sulfide minerals (Dale et al., 2009; Böning et al., 2020), the H_2_S produced during organoclastic sulfate reduction (Ferdelman et al., 1999) diffuses back from the sediment into the anoxic bottom waters (Brüchert et al., 2003; Ohde and Dadou, 2018) with severe effects for the coastal life (Currie et al., 2018). Toxic H_2_S escapes are then mitigated by the activity of planktonic sulfur-oxidizing bacteria that detoxify sulfide (Lavik et al., 2009; Callbeck et al., 2018; Crowe et al., 2018) while generating nitrite and NH_4_^+^ that can either augment nitrification with complete ammonia oxidation (comammox), or anammox to induce N_2_O emissions (Naqvi et al., 2010; Heiss and Fulweiler, 2016; Long et al., 2021). Trophic equilibrium in coastal waters is then rescued by the re-oxygenation of the SWI after the seasonal period of stratification. In austral winter 2018, water column and sediment samples suitable for microbiology analyses were obtained in the framework of an oceanographic sampling expedition to the BUS (Orsi et al., 2020b, 2022).

Here, water column geochemistry is combined with taxonomic, metagenomic and metatranscriptomic sequencing data from the water column, seafloor sediment, as well as water and sediment samples incubated for stable-isotope probing (SIP) to detail the taxonomy and metabolic activities of microbial populations involved in the enzymology and ecology of the nitrogen cycle (Lam and Kuypers, 2011; Martínez-Espinosa et al., 2011) under decreased stratification in Namibian shelf waters. The relative expression of the corresponding functional marker genes (Suter et al., 2021) and geochemical profiles for the water column and sediment (Siccha and Kucera, 2018; Ferdelman et al., 2021b, 2021a) show that active nitrification proceeds in two steps coupling ammonia and nitrite oxidation with respiratory nitrite reduction from oxic into dysoxic waters, and overall minor canonical denitrification in the quasi-absence of anammox (Long et al., 2021).

## Materials and Methods

### Sampling

The research expedition Meteor M148-2 to the Benguela upwelling, entitled EreBUS (i.e. Processes controlling the Emissions of gREenhouse gases from the Benguela Upwelling System) took place in 2018 from July 2^nd^ to 20^th^ (Ferdelman et al., 2019). The water column and underlying sediments were sampled at multiple sites on the shelf and then offshore (Weatherall et al., 2021) as the ship transited from Walvis Bay, Namibia to Las Palmas, Canary Islands (Fig. 1A). Water and sediment samples retrieved from the Namibian continental shelf (18.0 °S, 11.3 °E) were conditioned directly on board of the F/S Meteor vessel during the expedition (Orsi et al., 2020b, 2022). Water samples were retrieved using a Niskin rosette equipped with a Conductivity-Temperature-Depth system and a captor for light fluorescence reemitted by chlorophyll-a (chl-a) (Lavik et al., 2009; Siccha and Kucera, 2018) and water optical water quality variables (i.e. nepheloid turbidity unit) (Garaba et al., 2021). Water samples from the Niskin bottles were analyzed for dissolved nutrient concentrations (i.e. nitrate, nitrite, silica, phosphate) with a QuAAtro39 autoanalyzer (Seal Analytical) and fluorescence methods (i.e. NH_4_^+^) (Ferdelman et al., 2021b). At each site, 2 L of seawater were filtered via peristaltic pumping onto an in-line 0.2 μm polycarbonate filter. Replicate filters were then stored in sterile DNA/RNA clean 15 mL Falcon tubes and frozen immediately at −80 °C for further DNA and RNA analyses. A 30 cm-long gravity core was obtained from a water depth of 125 m at site 6, deploying a multicorer that yielded an intact SWI and the upper 30 cm of underlying sediment (Supplementary Fig. 1). The core was sectioned into 2 cm intervals, and sediments transferred into sterile DNA/RNA free 50 mL Falcon tubes and immediately frozen at −80 °C until DNA and RNA extractions (Orsi et al., 2020b, 2020a). Pore waters were extracted using MicroRhizon moisture samplers (Rhizon CSS, Rhizosphere research products) and analyzed for phosphate, nitrogen and sulfur species (Ferdelman et al., 2021a).

**Figure 1.**
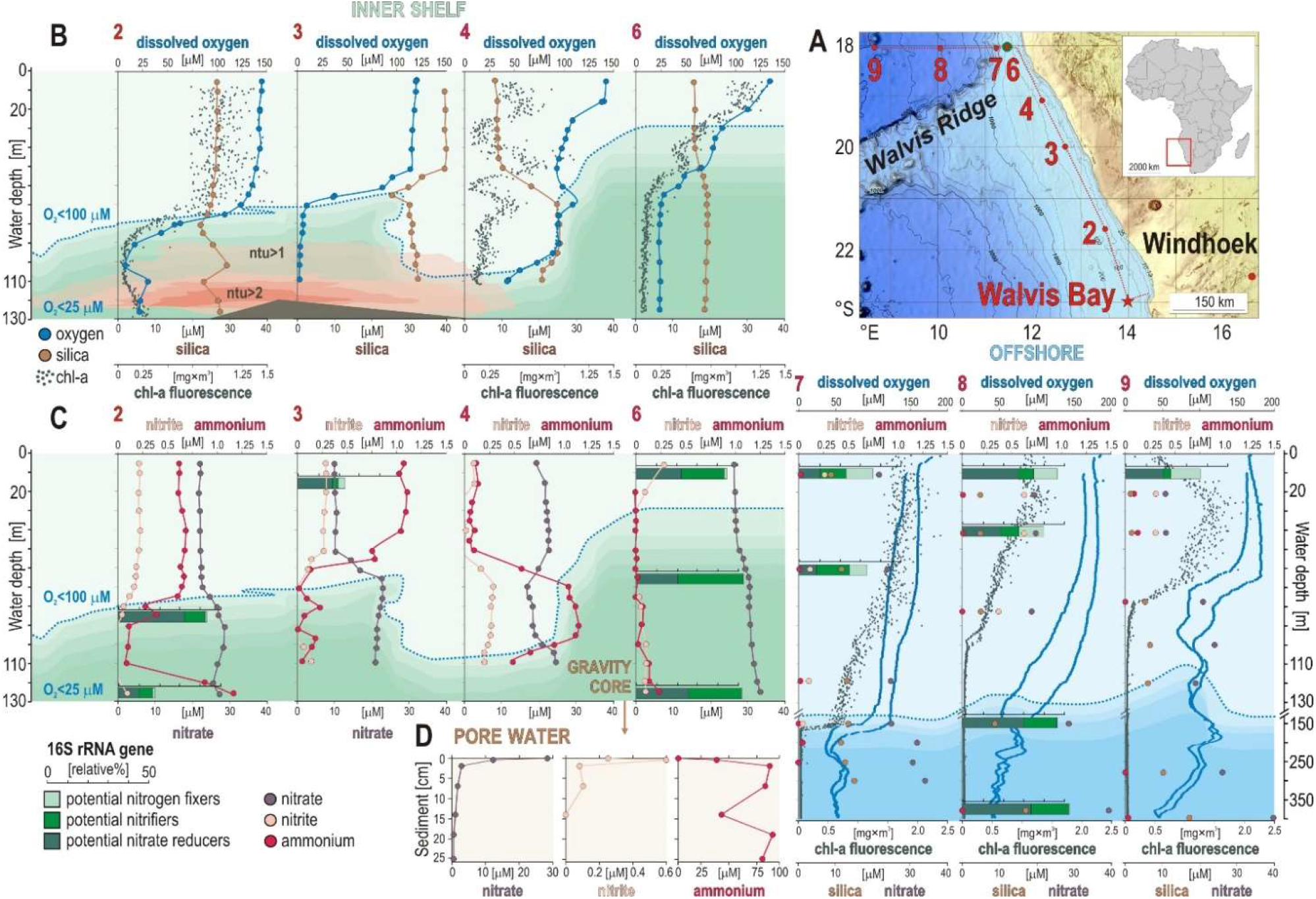
Sampling sites along the Namibian shelf, water column geochemical profiles, with relative abundance of microbial guilds involved in nitrogen cycling, and pore water geochemistry. (**A**) Bathymetric map of the Namibian shelf (data from Weatherall et al., 2021) displaying the different sites sampled during the EreBUS cruise 2018. (**B**) Concentration profiles for dissolved oxygen (blue) with an oxycline defined as <100 μM (dotted line), silica (brown), chlorophyll-*a* (green) and turbidity in nephelometric turbidity units (ntu) at each successive sampling site. (**C**) Concentration profiles for nitrite (pink), ammonia (red) and nitrate (purple) in the water column at each successive site, with relative abundances (% bar charts) of potential nitrogen fixers (light green), potential nitrifiers (green) and potential nitrate reducers (dark green). (**D**) Pore water geochemical profiles for nitrate (purple), nitrite (pink) and ammonium (red). Geochemical data are adapted from (Siccha and Kucera, 2018; Ferdelman et al., 2021b, 2021a; Garaba et al., 2021).

### Nucleic acid extractions

For size-fractionated filters, DNA was extracted by adding 850 mL of a sucrose ethylenediaminetetraacetic acid (EDTA) lysis buffer (0.75 M sucrose, 0.05 M Tris-Base, 0.02 M EDTA, 0.4 M NaCl, pH 9.0) and 100 mL of 10% sodium dodecyl sulfate to 2 mL bead-beating tubes containing the filters and 0.1 mm sterile glass beads, following a published protocol (Orsi et al., 2015; Vuillemin et al., 2022). After bead beating for 1 min and heating at 99 °C for 2 min, we added 25 mL of 20 mg × mL^−1^ proteinase K to the samples and incubated them at 55 °C overnight. DNA was extracted and purified from the lysate using the DNeasy Blood and Tissue Kit (Qiagen). The DNA from the sediments was extracted from 2 g using a sodium phosphate buffer and concentrated into 50 KDa amicon filters, as described in previous publications (Vuillemin et al., 2019, 2020b). DNA concentrations were quantified using a Qubit 3.0 fluorometer (Thermo Fisher Scientific).

RNA was extracted from either 2 g of sediment, or from filters, using the FastRNA Pro Soil-Direct Kit (MP Biomedicals) following the manufacturer’s instructions, with final elution of templates in 40 μL PCR water (Roche) as described previously (Vuillemin et al., 2020b, 2020a). Extraction from the filters was processed using 4 mL of RNA lysing solution together with silica glass beads from two Lysing Matrix E tubes, and homogenized using the FASTprep 5-G homogenizer (all MP Biomedicals). In order to maximize recovery of the RNA pellet, we added 4 μL glycogen at 1 μg × mL^−1^ prior to the 30 min isopropanol precipitation. All RNA samples were extracted in a HEPA-filtered laminar flow hood and set of pipets exclusively dedicated to RNA samples. Before and after each extraction, all surfaces were treated with RNAse-Zap and exposed to UV light for 30 min. Pipets were systematically autoclaved after use.

### Stable-isotope probing incubations, 16S rRNA gene quantification

Water samples from 10 mwd and 125 mwd at site 6 were selected for SIP incubations and amended with ^13^C-labeled diatom necromass produced from a culture of *Chaetocerous socialis* (Norwegian Culture Collection strain K1676), as previously described (Orsi et al., 2022). The concentrated mixture of dead diatom exopolysaccharides (dEPS) was used as ^13^C-labeled substrate (^13^C enrichment of >50%) for tracing *in vitro* heterotrophic microbial activities by amending ^13^C-labeled OM at a final concentration of 200 μg × g^−1^ (1–3% of the *in situ* carbon concentrations). Sediments from 28 cm below the seafloor (cmbsf) at site 6 were selected for SIP incubations amended with ^13^C-bicarbonate (DIC). Flasks were incubated in triplicate in the dark for 18 hours (dEPS) and 10 days (DIC), respectively and frozen to −80°C to terminate the incubations. DNA was extracted from the incubation slurries in the home lab, as described above. DNA extracts were fractionated into 15 pools via density gradient according to published SIP protocols (Hungate et al., 2015; Coskun et al., 2022), resuspended into 30 μL molecular grade (DEPC-treated) water and quantified using a Qubit 3.0 fluorometer.

To determine shifts in the peak buoyant density of DNA of the incubations, qPCR assays targeting the V4 hypervariable region of 16S rRNA genes were carried out on the 15 density fractions. DNA templates were used in qPCR amplifications with updated 16S rRNA gene primer pair 515F (5’-GTG YCA GCM GCC GCG GTA A −3’) with 806R (5’-GGA CTA CNV GGG TWT CTA AT −3’) to increase the coverage of Archaea and marine clades (Parada et al., 2016) and run as previously described (Coskun et al., 2019, 2022). The reaction efficiencies in all qPCR assays were between 90% and 110%, with an r^2^ of 0.98. Gene copies were normalized to the wet weight of the sediment and volume of water filtered. The ^13^C-labeled fractions with highest gene copy numbers were selected and pooled for metagenomic library preparation (Supplementary Fig. 2).

### Library preparation

Amplicons of tagged partial 16S rRNA gene primers were run on 1.5% agarose gels, the bands excised and purified with the QIA quick Gel Extraction Kit (QIAGEN) and the final eluted DNA quantified with the Qubit dsDNA HS Assay Kit (Thermo Fisher Scientific). Purified PCR amplicons containing unique barcodes from each sample were diluted to 1 nM solutions and pooled.

For metagenomes, initial DNA extracts were diluted to DNA concentrations of 0.2 ng × μL^−1^ and used in metagenomic library preparations with the Nextera XT DNA Library Prep Kit (Illumina Inc.). For metatranscriptomes, DNAse treatment, synthesis of complementary DNA and library construction using specific barcodes were obtained from 10 μL of RNA templates by processing the Trio RNA-Seq kit protocol (NuGEN Technologies). Because the Trio RNAseq Ovation kit (NuGEN technologies) is biased against molecules with secondary structure such as ribosomal RNA and preferentially amplifies messenger RNA, performing a ribosomal RNA (rRNA) depletion step was not necessary. All libraries were quantified on an Agilent 2100 Bioanalyzer System, using the High Sensitivity DNA reagents and DNA chips (Agilent Genomics), diluted to 1 nM, and pooled according to the MiniSeq System Denature and Dilute Libraries Guide from Illumina. 500 μL library (1.8 pM) with 8 μL denatured PhiX control were sequenced in two separate runs using an paired-end Mid Output Kit (300-cycles) on the Illumina MiniSeq platform (Pichler et al., 2018).

### Assembly and analysis

Demultiplexing and base calling were performed using bcl2fastq Conversion Software v. 2.18 (Illumina, Inc.). MiniSeq read trimming and assembly, OTU picking and clustering at 97% sequence identity were done using USEARCH (Edgar, 2013). Taxonomic assignments of 16S rRNA genes were generated by QIIME, version 1.9.1 (Caporaso et al., 2010), using the implemented BLAST method against the SILVA rRNA gene database, release 138 (Quast et al., 2013). All OTUs containing <10 sequences and which had no BLASTn hit were removed. Reads passing this quality control were then normalized by percentage of total sequencing depth per sample.

For metagenomic and metatranscriptomic MiniSeq reads, quality control, *de novo* assembly and open reading frames (ORFs) searches were performed as described previously (Ortega-Arbulú et al., 2019). Reads were trimmed and paired-end reads assembled into contigs, using CLC Genomics Workbench 9.5.4 for a minimum contig length of 300 nucleotides. Reads were then mapped to the contigs using the following parameters (mismatch penalty = 3, insertion penalty = 3, deletion penalty = 3, minimum alignment length = 50% of read length, minimum percent identity = 95%). Coverage values were obtained from the number of reads mapped to a contig divided by its length (i.e. average coverage). Only contigs with an average coverage >5 were selected for ORF searches and downstream analysis. This protocol does not assemble ribosomal RNA, and thus transcript results are only discussed in terms of messenger RNA. Protein encoding genes and ORFs were extracted using FragGeneScan v. 1.30 (Rho et al., 2010). For 16S rRNA gene amplicon datasets, each sample was sequenced to an average depth of 22,876 sequences per sample. Metagenomes were sequenced to an average depth of 6.3 million reads per sample. Metatranscriptomes spanning water column and seafloor habitats (n = 27) were sequenced with an average depth of 5.9 million reads, and after *de novo* assembly an average of 17,943 contigs per sample could be assembled. For the metagenomes prepared from the ^13^C-labeled SIP fractions, libraries were sequenced to an average depth of 6.6 million reads (Supplementary Table 1 and 2).

Taxonomic identifications were integrated with the functional annotations, performing BLASTp and BLASTx searches of ORFs against a large aggregated genome database of predicted proteins using the DIAMOND protein aligner version 0.9.24 (Buchfink et al., 2015). The aggregated genome database of predicted proteins includes the SEED (www.theseed.org) and NCBI RefSeq databases updated with all predicted proteins from recently described high-quality draft subsurface metagenomic assembled genomes (MAGs) and single-cell assembled genomes (SAGs) from the NCBI protein database. The coverage of total annotated protein-encoding ORFs detected, as opposed to the number of reads mapping per kilobase per ORF (for example, RPKM), was selected to reduce potential bias from small numbers of “housekeeping” genes with potentially higher expression levels. This approach allows to assign ORFs from the metagenomic and metatranscriptomic data to higher-level taxonomic groups, and thereby to draw environmental conclusions about their specific metabolic traits and activities (Breitwieser et al., 2019). The complete bioinformatics pipeline has been previously published (Orsi et al., 2020b, 2022). Statistical analyses were performed using RStudio v. 3.3.3 with the Bioconductor package (Huber et al., 2015). All scripts and codes used to produce sequence analyses have been posted on GitHub with a link to the instructions on how to conduct the scripts (github.com/williamorsi/MetaProt-database).

### Phylogeny of functional genes

All 16S rRNA gene amplicon sequences were aligned with SINA online v. 1.2.11 (Pruesse et al., 2007) and inserted in the SILVA 16S rRNA SSU NR99 reference database tree, release 138 (Quast et al., 2013), using the maximum parsimony algorithm without allowing changes of tree typology. Partial OTU sequences closely affiliated with known nitrifiers were selected with their environmental references and plotted in separate archaeal and bacterial Maximum Likelihood RAxML phylogenetic trees, using rapid bootstrap analysis and selecting the best trees among 100 replicates using ARB (Ludwig et al., 2004).

Phylogenetic analyses of the predicted ammonia monooxygenase (*amoA*) and copper-containing nitrite reductase (*nirK*) gene proteins were performed for all the corresponding annotated taxa in the metagenomes and metatranscriptomes, using 185 and 133 aligned amino acid sites, respectively (Garbeva et al., 2007; Alves et al., 2018). For each of the two marker gene phylogenies (amoA, *nirK*), all ORFs annotated to those genes from the bioinformatics pipeline were aligned against their top two BLASTp hits in the NCBI-nr and SEED databases using MUSCLE (Edgar, 2004). Conserved regions of the alignments were selected and phylogenetic analyses of the predicted proteins were performed in SeaView version 4.7 (Gouy et al., 2010) using RAxML (Stamatakis, 2014) with BLOSUM62 as the evolutionary model and 100 bootstrap replicates.

## Results

### Water column geochemistry, taxonomic assemblages across sampling locations

Consistent with prior studies (Kuypers et al., 2005; Lavik et al., 2009), an oxycline was observed between 50 and 95 mwd that exhibited O_2_ concentrations spanning 100-40 μM O_2_ (Fig. 1A). Below 65 to 95 mwd, an OMZ defined as having <60 μM O_2_ (Wright et al., 2012) was detected at all sites sampled along the shelf (Fig. 1A). The profiles measured for chl-a fluorescence (Siccha and Kucera, 2018) display maximum concentrations (i.e. 1.25 mg × m^3^) in the surface ocean at site 6, with a general trend decreasing with water depth that runs in parallel to the oxygen profiles (Fig. 1B). Silica covarying with *chl-a* concentrations may reflect productivity by siliceous plankton like diatoms (Inthorn et al., 2006b), otherwise sediment suspension in nepheloid layers (Inthorn et al., 2006a). Turbidity in the water column, measured as nephelometric turbidity units (ntu), is detectable (i.e. ntu >1) in the shelf bottom waters between site 2 and site 4 (Fig. 1B). Together with the *chl-a* fluorescence profile at site 4, these data indicate the presence of water eddies along the continental shelf with suspension of seafloor sediments into the water column (Fig. 1B). This is typical of the winter months on the Namibian coast that present more well-mixed water conditions (Brüchert et al., 2009; Ohde and Dadou, 2018; Callbeck et al., 2021).

Concomitant trends in NH_4_^+^, nitrite and nitrate profiles can indicate nitrification, denitrification and anammox processes (Fig. 1C). Based on measured concentrations, there appears to be different processes at sites 2 and 3 compared to sites 4 and 6 where phosphate concentrations respectively decrease and increase in the OMZ (Supplementary Fig. 3). Site 4 displays geochemical features that are indicative of denitrification with some nitrification processes above and within the OMZ, i.e. respectively a decrease in nitrate and NH_4_^+^ and increase in nitrite concentrations (Fig. 1C), whereas sites 2 and 3 display decreasing concentrations in NH_4_^+^ and nitrite in the OMZ while nitrate increases (Fig. 1C). Such concomitant decrease in nitrite and NH_4_^+^ could also be indicative of anammox processes in the OMZ at sites 2 and 3, and above the OMZ at sites 4 and 6.

However, geochemical profiles in the water column cannot be attributed to *in situ* microbial processes of nitrogen cycling only, but also inform on nutrient sources and transport. NH_4_^+^ concentrations in bottom waters at site 2 reveal that benthic releases locally occur on the shelf with transport in the benthic boundary layer along the shore (Brüchert et al., 2009; Füssel et al., 2012; Schunck et al., 2013; Neumann et al., 2016; Ohde and Dadou, 2018; Hanz et al., 2019). In the sediment underlying bottom waters of the OMZ at site 6, pore water chemical analysis indicates that nitrate and nitrite were consumed quickly at the sediment surface, followed by an increased accumulation of NH_4_^+^ with depth (Fig. 1D). The sediments exhibit a redox gradient spanning dysoxic conditions (~25 μM O_2_) at the seafloor surface to sulfidic conditions at 30 cmbsf (Supplementary Fig. 4). In contrast to site 2 (Fig. 1C), NH_4_^+^ does not diffuse across the SWI at site 6 (Fig. 1D).

In the water column, sequencing results of 16S rRNA genes show that the microbial assemblages mostly consist of Cyanobacteria, Bacteroidota (i.e. Bacteroidia, Chlorobia, Ignavibacteria), Alpha- and Gammaproteobacteria. The class Nitrososphaeria (former phylum Thaumarchaeota), which represents 10% of the total microbial composition in surface waters at all sites, sums up to 30% of total 16S rRNA genes in the OMZ waters at site 6. In the sediment, the 16S rRNA gene assemblage shows a clear predominance of Chloroflexi, complemented with mostly Desulfobacterota and Gammaproteobacteria, and only 2% on average of Archaea. The copy number of 16S rRNA genes obtained via qPCR assays is highest at the SWI (i.e. log10=8) and remains relatively constant (i.e. log10=6-7) with sediment depth. The corresponding non-metric multidimensional scaling (NMDS) analysis based on 12,200 OTUs clearly indicates a significant shift in the community composition from the surface ocean across the OMZ waters down to the SWI, and further into the sediment (Supplementary Fig. 5).

Analysis of the metagenomes provided a way to cross-check taxonomic assignments based on 16S rRNA primers. Comparison between the two showed a similar archaeal distribution across water column samples, whereas ORFs assigned to Proteobacteria were somehow increased in the metagenomes. In the sediment, the number of ORFs assigned to Gammaproteobacteria apparently increased at the expenses of those of Chloroflexi. In the metatranscriptomes, protein-encoding ORFs were by far more assigned to Desulfobacterota along the shelf waters (sites 2 and 4), whereas those from northward sampling sites mostly indicated Bacteroidota, Alpha- and Gammaproteobacteria, and Nitrososphaeria as the main metabolically active phyla. In the sediment, Firmicutes, Desulfobacterota and Gammaproteobacteria represented about 50% of the expressed ORFs. The NMDS analysis based on all annotated protein-encoding ORFs obtained from our metagenomes and metatranscriptomes clearly separates all of the metagenome and metatranscriptome samples, plotting samples at different sampling locations spanning from the surface ocean across OMZ waters to sediment core top to bottom (Supplementary Fig. 5).

### Diversity and abundance of nitrogen cycling populations

Relative abundances based on 16S rRNA genes were established for sub-populations putatively involved in nitrogen fixation (among Actinobacteriota, Cyanobacteria, Firmicutes and Alphaproteobacteria), nitrifiers (among Nitrososphaeria, Nitrospinota, Nitrospirota and Gammaproteobacteria) and nitrate reducers (among Bacteroidota, Firmicutes, Nitrospirota, Alphaproteobacteria, Desulfobacterota and Gammaproteobacteria) to determine the main taxa driving the nitrogen cycle across all sites sampled along the shelf (Fig. 2).

**Figure 2.**
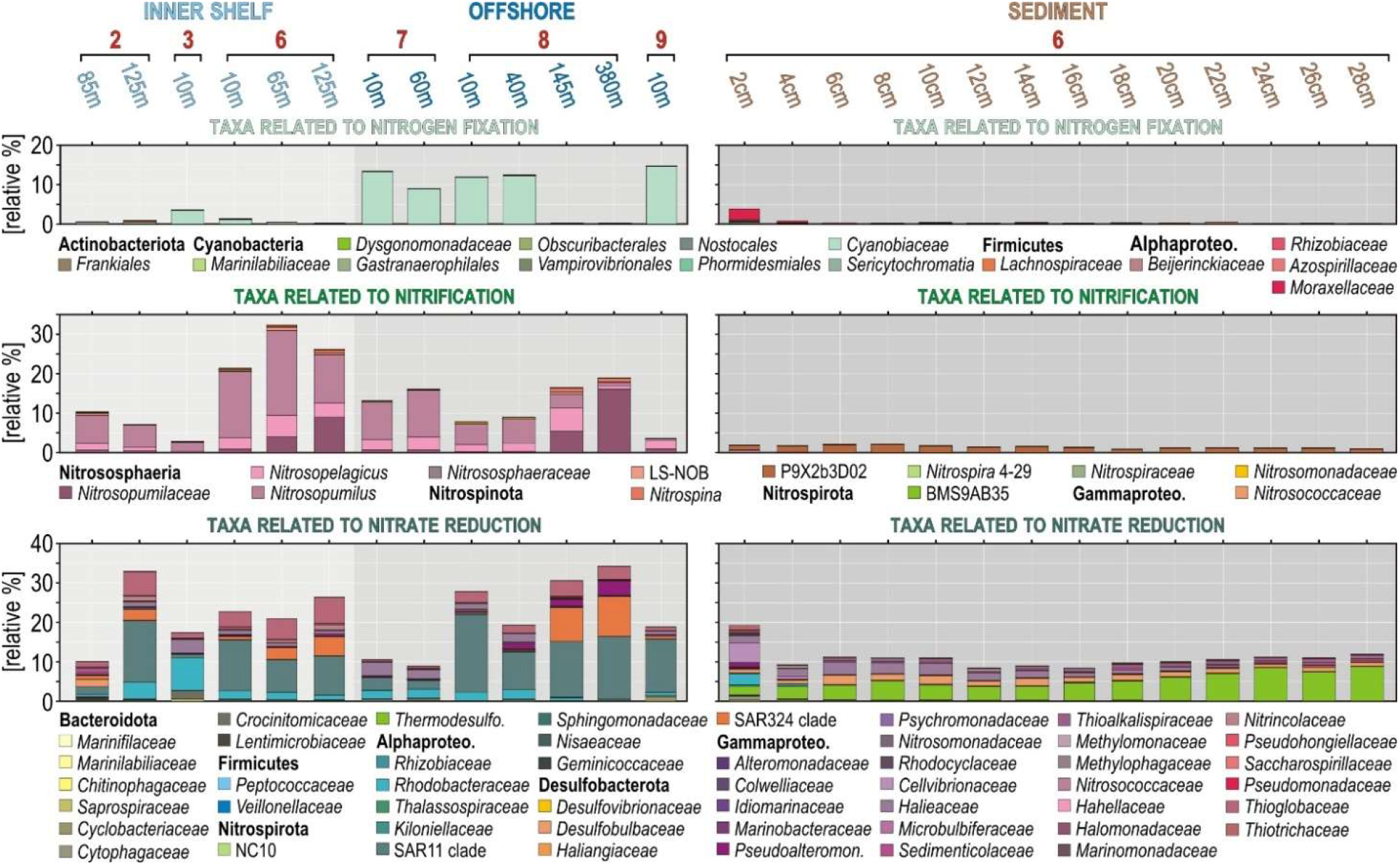
Quantification and taxonomy of 16S rRNA genes grouped into functional guilds for potential nitrogen fixers, nitrifiers and nitrate reducers. (**Top to bottom**) Relative abundances and taxonomy of taxa potentially involved in nitrogen fixation (top), nitrification (middle) and nitrate reduction (bottom) in the water column across successive sampling sites (left) and in the sediment (right).

Taxa involved in nitrogen fixation in the ocean surface are more abundant at offshore stations (site 7 and 8), corresponding predominantly to cyanobacterial populations, whereas diazotrophic Alphaproteobacteria were exclusively identified at the SWI and in the uppermost sediment (Fig. 2). Populations potentially carrying out nitrification increase from the surface ocean into the OMZ waters, especially at site 6 and 7 where the top part of the oxycline is located at shallower water depths (i.e. 20-40 mwd).

Ammonia-oxidizing archaea (AOA) are the most abundant nitrifiers in the water column, constituting up to 30% of the entire 16S rRNA gene assemblage in OMZ waters. The detailed phylogenetic analysis indicates the presence of *Candidatus* (Ca.) Nitrosopelagicus and *Ca*. Nitrosopumilus in the water column, and *Ca*. Nitrososphaera in the sediment (Fig. 3A), respectively separating planktonic (27 OTUs) and benthic (9 OTUs) populations. Nitrite-oxidizing bacteria (NOB) among Nitrospinota are present at very low 16S rRNA relative abundances (Fig. 2B), but are taxonomically diverse with 60 OTUs distributed among candidate clade LS-NOB and *Nitrospinaceae* in the water column, and candidate clade P9X2b3D02 in the sediment (Fig. 3B). Other NOB and potential ammonia-oxidizing bacteria (AOB) are not abundant either (Fig. 2), including diverse taxa among the Nitrospirota (8 OTUs) and Gammaproteobacteria (20 OTUs), respectively affiliated with *Nitrospira*, *Nitrosomonas*, and *Nitrosococcus* (Fig. 3B).

**Figure 3.**
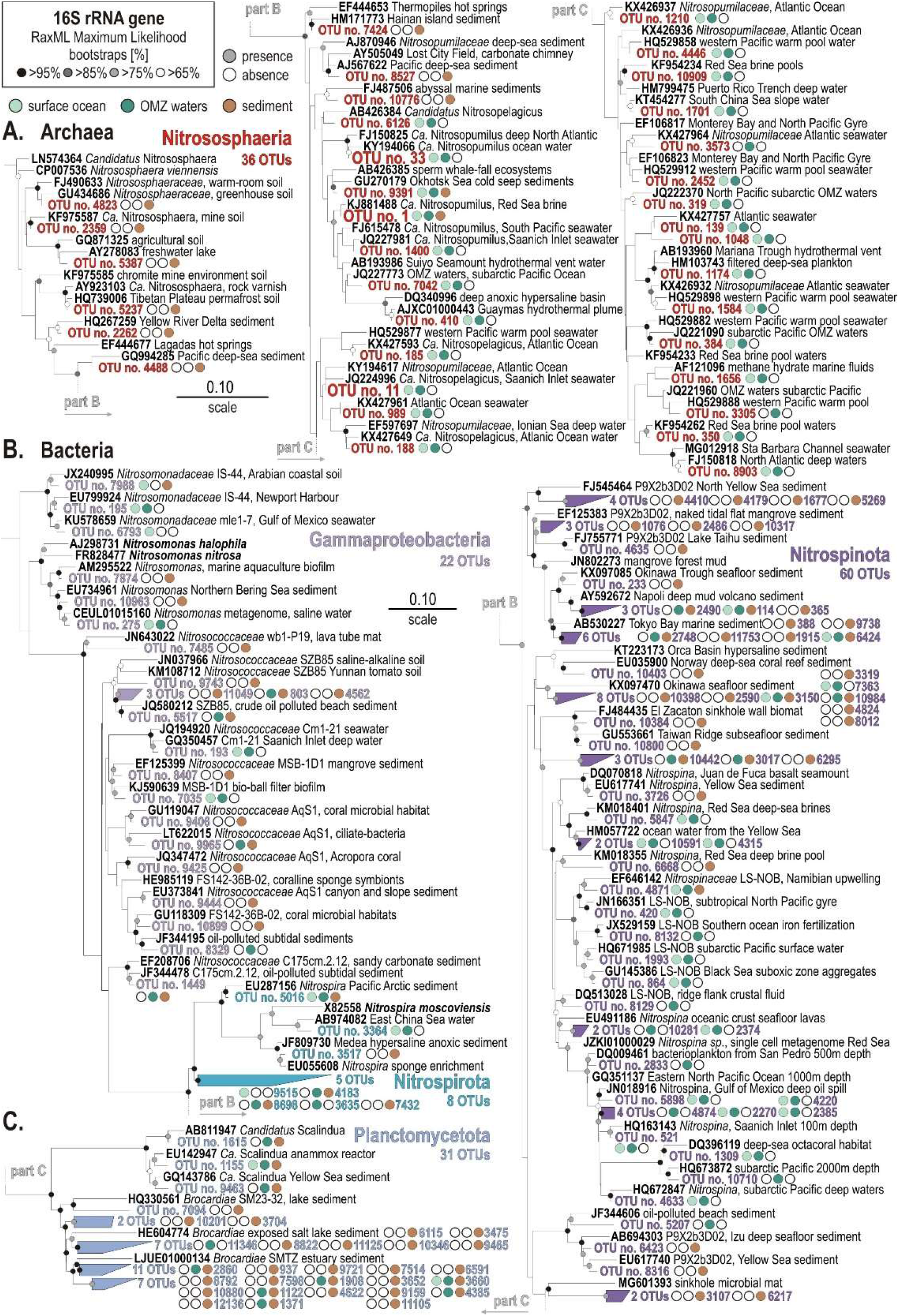
Phylogenetic analysis of partial 16S rRNA gene amplicons for ammonia-oxidizing archaea (A), ammonia- and nitrite-oxidizing bacteria (B), and anammox bacteria (C). RAxML Maximum Likelihood archaeal and bacterial trees selected among 100 replicates for all partial 16S rRNA transcripts (V4 hypervariable region). Presence/absence of a 16S rRNA gene with the surface ocean (light green), OMZ waters (dark green) and sediment (brown) is signified by full versus empty circles. Bold types signify accession numbers and cultivated species, whereas regular font indicates the sequence isolation sources.

Nitrate-reducing bacteria potentially involved in denitrification and DNRA represent up to 30% of the total 16S rRNA gene assemblages in OMZ deep waters. Predominant taxa in the water column are affiliated with the alphaproteobacterial family *Rhodobacteraceae* and *Thalassospiraceae*, (formerly deltaproteobacterial) clade SAR324, and gammaproteobacterial family *Pseudoalteromonadaceae* and *Thioglobaceae* (Fig. 2). The SWI and surface sediment are apparently colonized by *Rhodobacteraceae, Cellvibironaceae, Halieaceae* and *Thiotrichaceae*, along with *Thermodesulfovibrionaceae* and *Desulfobulbaceae*, which are also known to include sulfur-oxidizing and sulfate-reducing bacteria potentially carrying out DNRA. Consistent with the depletion of pore water nitrate, the relative abundances of these taxa decrease in shallow sediment as the sediment becomes sulfidic (Supplementary Fig. 4).

We also searched for 16S rRNA gene sequences affiliated with Planctomycetota known to perform anammox. We identified 31 OTUs that closely match *Ca*. Scalindua and *Ca*. Brocardia, among which only 10 OTUs are derived from OMZ water samples whereas 21 OTUs are solely present in the sediment (Fig. 3C). However, these 31 OTUs are minor in terms of relative abundances.

### Stable-isotope probing incubations and ^13^C-labeled taxa

After 18 hours of incubation in the dark at 10 °C, the ^13^C-dEPS incubations showed ^13^C-labeling of 16S rRNA genes, defined by a shift in peak DNA buoyant density, in the surface ocean (10 mwd) and OMZ (125 mwd), resulting in “isotopically heavier” (i.e. ^13^C-enriched DNA) 16S rRNA genes compared to the controls that received the unlabeled substrate (Supplementary Fig. 2). Similarly after 10 days of incubation in the dark at 10 °C, the ^13^C-bicarbonate incubations amended with subseafloor sediment (Orsi et al., 2020b, 2022) showed an increased buoyant density of 16S rRNA genes compared to the unlabeled controls. This indicates that ^13^C-labeling of microbes synthesizing new DNA had occurred, which had assimilated the added ^13^C-dEPS and ^13^C-DIC into their biomass.

Metagenomic sequencing of these “isotopically heavy DNA” SIP fractions shows that most of the planktonic bacteria that had assimilated the added ^13^C-dEPS substrate are affiliated with Bacteroidota, both within the 10 mwd and 125 mwd (OMZ) incubations. Additionally, Nitrososphaeria increase in relative abundance at the highest CsCl densities, indicating that putative heterotrophic Nitrososphaeria (Kitzinger et al., 2019; Aylward and Santoro, 2020) had also potentially assimilated ^13^C-dEPS in the SIP incubations. Taxonomic affiliations of the ORFs in the sediment heavy SIP metagenomes reveal ^13^C assimilation mainly by the Desulfobacterota, Gammaproteobacteria, Chloroflexi and Actinobacteria (Supplementary Fig. 2).

### Potential and expression of functional marker genes involved in nitrogen cycling

Here, the relative % of total prokaryotic ORFs identified in the metatranscriptomes is compared to those present in the metagenomes in order to assess levels of metabolic expression in the enzymology of nitrogen cycling. The focus is set on ORFs encoding genes involved in nitrogen regulatory metabolism, diazotrophy, nitrification, denitrification, and simple organic nitrogen compound oxidases, reductases and transporters (Lam and Kuypers, 2011; Martínez-Espinosa et al., 2011; Daims et al., 2016). The presence and expression of ORFs with correspondence to P_II_ proteins, ammonia lyases and ammonia ligases were used to trace nitrogen regulatory metabolism. Diazotrophy and nitrification were investigated via protein-encoding genes involved in nitrogen fixation (*nif*), ammonium monooxygenase (amo), nitrite oxidoreductase (nxr), nitronate monooxygenase also known as 2-nitropropane dioxygenase (nmo), along with ureases, nitrilases and cyanases. ORFs with correspondence to gene subunits encoding the respiratory nitrate reductase (*nar*), periplasmic nitrate reductase (*nap*), nitrite reductase (nir), nitric oxide reductase (nor), nitrous oxide reductase (*nos*), and ammonia-forming nitrite reductase (*nrf*) were sorted to trace denitrification and DNRA. For organoclastic processes in link to denitrification, genes encoding nitroreductases (ntr), nitrilases (nit) and formate dehydrogenase (*fhn*), as well as for nitrate/nitrite, NH_4_^+^ and urea transporters (Fig. 4) were investigated. In addition, for hydrazine hydrolase and hydrazine oxidoreductase (hzo) which is involved in anammox processes (Karlsson et al., 2009), a single ORF was detected in the metagenome from 50 mwd at site 4.

**Figure 4.**
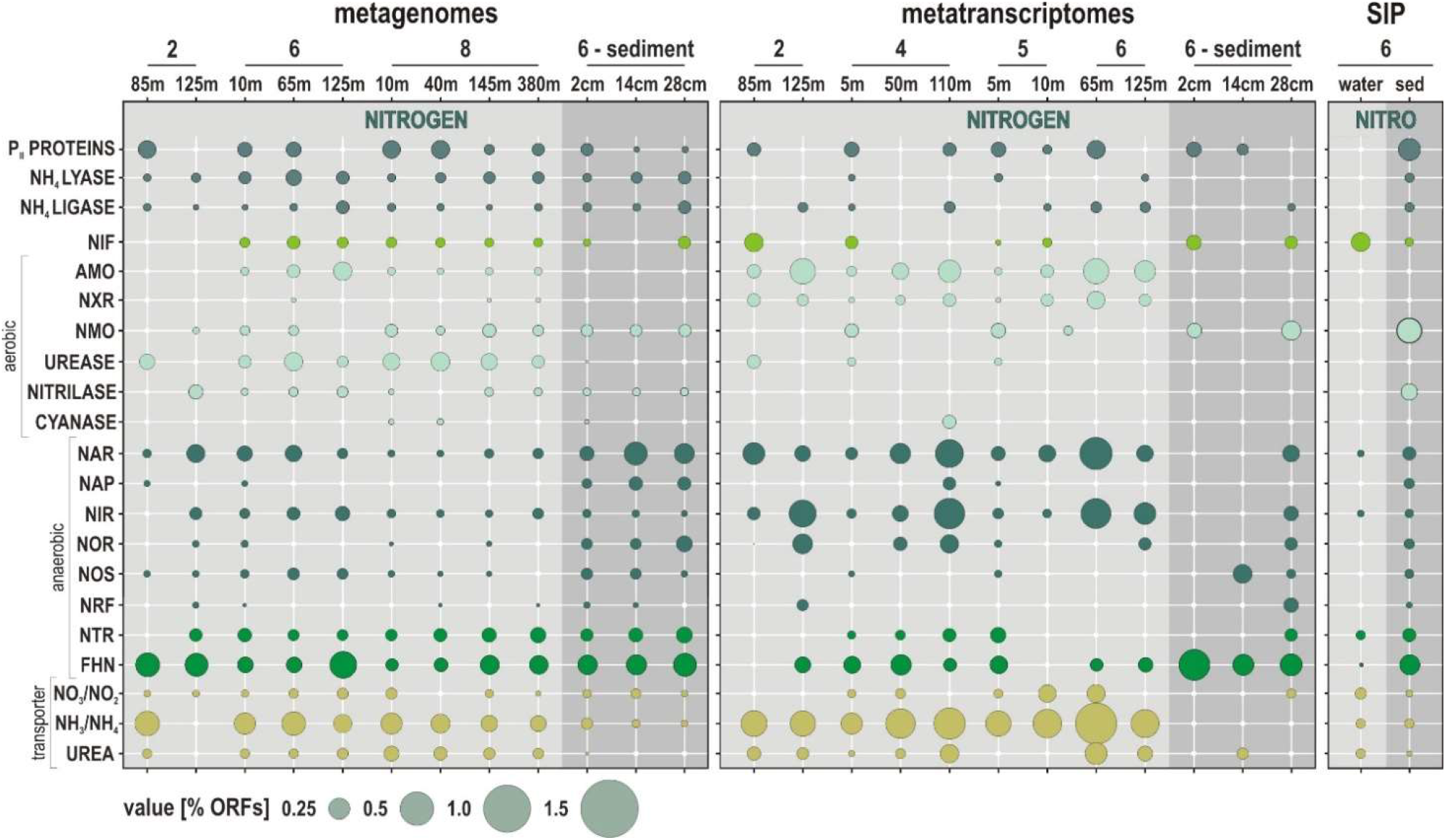
Metabolic potential and activities related to nitrogen cycling along the Namibian coast. Bubble plot showing the relative abundances of metabolic functions [% total ORFs] in the metagenomes, metatranscriptomes, and stable isotope probing (SIP) sediment incubations (left to right) assigned to functional marker genes involved in nitrogen cycling. *Abbreviations:* P_II_ PROTEINS: nitrogen regulatory proteins / NIF: nitrogen fixation / AMO: ammonium monooxygenase / NXR: nitrite oxidoreductase / NMO: nitronate monooxygenase / NAR: nitrate reductase / NAP: periplasmic nitrate reductase / NIR: copper-containing nitrite reductase / NOR: nitric oxide reductase / NOS: nitrous oxide reductase / NRF: ammonia-forming cytochrome nitrite reductase / NTR: nitroreductases / FHN: formate dehydrogenase.

The relative expression of ORFs attributed to P_II_ proteins, NH_4_^+^ lyases and *nif* genes is indicative of related metabolic activity limited to surface waters, whereas those of NH_4_^+^ ligases tend to increase into OMZ waters. ORFs related to *amo* and *nxr* genes, with concomitant nitrite and NH_4_^+^ transporters, are expressed at relatively high % in the water column at all sites (Fig. 4). The number of these ORFs shows a clear tendency to increase towards the oxycline and into OMZ waters, whereas they are below detection in the anoxic sediment. This indicates that nitrification processes are active and even increase in the vicinity of the oxycline. ORFs related to ureases and urea transporters are mostly expressed in oxic waters along the shelf. Expression of ORFs encoding nitrilases, cyanases and *nmo* genes is minor and only detectable in the surface ocean and oxic waters (Fig. 4).

The relative % of expressed ORFs assigned to *nar* and *nir* genes increases from surface into OMZ waters at all sites (Fig. 4). The relative % of ORFs assigned to *nor* genes follows the same trend, with expression mostly detected in OMZ waters. In comparison, expression of ORFs assigned to *nos* genes is minor and limited to surface waters. Although *nos* genes are usually considered to be unique to denitrifying bacteria (Scala and Kerkhof, 1999), their detection currently restricted to the surface ocean may indicate aerobic consumption of N_2_O gas emissions (Sun et al., 2017). Ammonification mediated by *nrf* gene transcription is only expressed in bottom waters on the shelf and in the sediment. Expressed ORFs assigned to nitroreductases were only detected at site 4, where eddies promote water column oxygenation and sediment suspension along the shore (Inthorn et al., 2006a). ORFs assigned to *fhn* genes, which is part of the respiratory chain of denitrification as well as of the acetogenic W-L pathway (Einsle and Kroneck, 2004), consistently increase from ocean surface into OMZ waters and underlying sediment (Fig. 4). At all sites, transcripts encoding the *amo*, *nxr* and *nir* genes and NH_4_^+^ transporters display the highest expression levels (calculated as % of total reads) (Fig. 5A), with taxonomic assignments demonstrating that Nitrosophaeria and Nitrospinota couple nitrification with denitrification (Lawton et al., 2013) in coastal OMZ waters (Fig. 5B).

**Figure 5.**
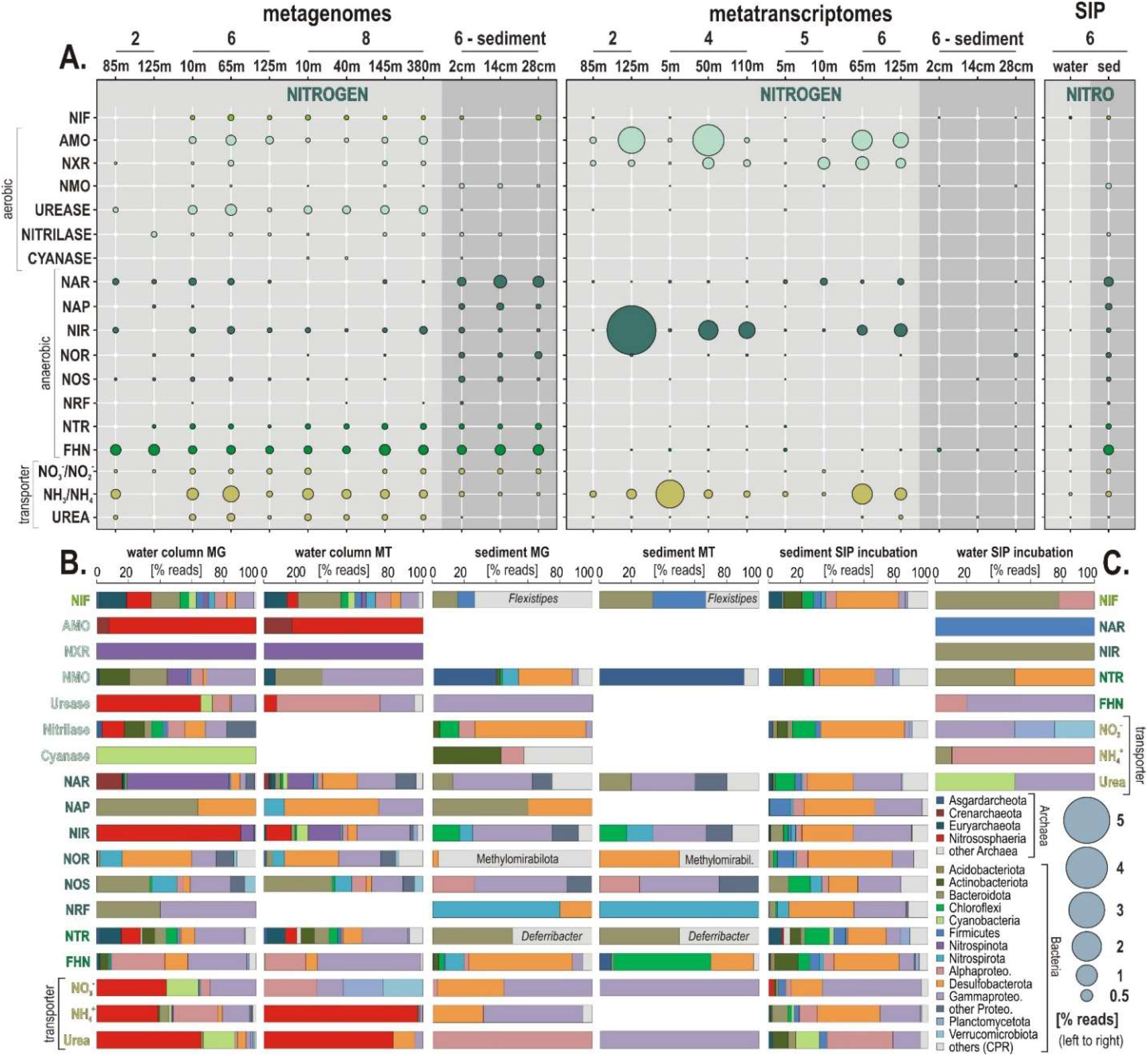
Metabolic functions and activities related to nitrogen cycling in the water column, sediment and SIP incubations, and the corresponding taxonomic assignments at the phylum level. **(A)** Bubble plot showing the relative potential and expression of metabolic functions [% total reads] assigned to nitrogen cycling in the metagenomes, metatranscriptomes, stable isotope probing (SIP) water and sediment incubations (left to right). (**B**) Taxonomic bar charts [% reads] for the corresponding functional marker genes related to nitrogen cycling at the phylum level in the metagenomes (MG), metatranscriptomes (MT) and anaerobic SIP incubations with sediment and ^13^C-labeled bicarbonate (DIC). The last column on the left (**C**) displays taxonomy of functional marker genes identified in aerobic incubations with water and ^13^C-labeled diatom mixture (dEPS). *Abbreviations:* NIF: nitrogen fixation / AMO: ammonium monooxygenase / NXR: nitrite oxidoreductase / NMO: nitronate monooxygenase / NAR: respiratory nitrate reductase / NAP: periplasmic nitrate reductase / NIR: copper-containing nitrite reductase / NOR: nitric oxide reductase / NOS: nitrous oxide reductase / NRF: ammonia-forming cytochrome nitrite reductase NTR: nitroreductases / FHN: formate dehydrogenase.

The ^13^C-labeled functional marker genes related to nitrogen cycling in SIP metagenomes obtained from water incubations reveal concomitant *in vitro* processes of nitrogen fixation (*nif*), denitrification (*nar, nir*), degradation (*ntr, fhn*) and uptake of nitrogen compounds (NO_2_^−^, NH_4_^+^, urea transporters) from ^13^C-dEPS, with denitrification initiating within 18 hours (Fig. 4). In SIP metagenomes from sediment incubations, the ^13^C-labeled functional marker genes are indicative of the complete denitrification pathway along with reduction of organic nitriles and nitronates (nitrilases, nmo). Because it is hard to discriminate ORFs genuinely occurring by isotopic enrichment in the “heavy” DNA fraction from those with high GC content (Coskun et al., 2022) and because the isotopically “light” DNA fractions were not sequenced, it may well be that the ^13^C-labeled ORFs with low GC content are not represented in the analysis.

### Taxonomic assignment of functional marker genes, phylogenetic analysis

To identify the main microbial constituents involved in nitrogen cycling across the coastal OMZ, the taxonomic assignment of the ORFs corresponding to the aforementioned functional markers genes were summed up separately for the metagenomes, metatranscriptomes and SIP incubations (Fig. 5), according to the isolation source (i.e. water column or sediment).

Briefly, populations with metabolic potential to fix nitrogen are taxonomically diverse (e.g. Bacteroidota, Cyanobacteria, Alphaproteobacteria), but their corresponding expression levels of *nif* proteins are very low (Figs. 5A–5B). Nitrosophaeria are clearly the main actors in active transcription of *amo* and *nir* genes, nitroreductases, ureases and urea transporters in the water column. Transcription levels of nxr, nar, and *nir* genes by Nitrospinota are relatively high as well (Figs. 5A–5B). Phyla actively encoding the complete denitrification pathway (i.e. nar, *nap, nir, nor, nos* genes) in the water column include Nitrospirota, Desulfobacterota and Gammaproteobacteria (Fig. 5B). Although these processes are minor in the sediment (Figs. 4 and 5A), phyla that express the related ORFs are mostly identified as Proteobacteria, and accessorily as Nitrospirota and Bacteroidota (Fig. 5B). Expression of *fhn* genes by Chloroflexi was considered indicative of acetogenesis (Vuillemin et al., 2020a).

The phylogenetic analyses of *amoA* and *nirK* gene sequences confirm that Nitrosophaeria (Fig. 6A) and Nitrospinota (Fig. 6B) are the main drivers of nitrifier denitrification (Fernandez and Farias, 2012; Lawton et al., 2013; Wrage-Mönnig et al., 2018). Assignments of *amoA* genes (Fig. 6A) show that only AOA among the class Nitrosophaeria are metabolically active, whereas AOB such as *Nitrosomonas* or *Nitrosococcus* were not identified. Canonical denitrifiers actively expressing *nirK* genes are assigned to Alpha- and Gammaproteobacteria (Fig. 6B).

**Figure 6.**
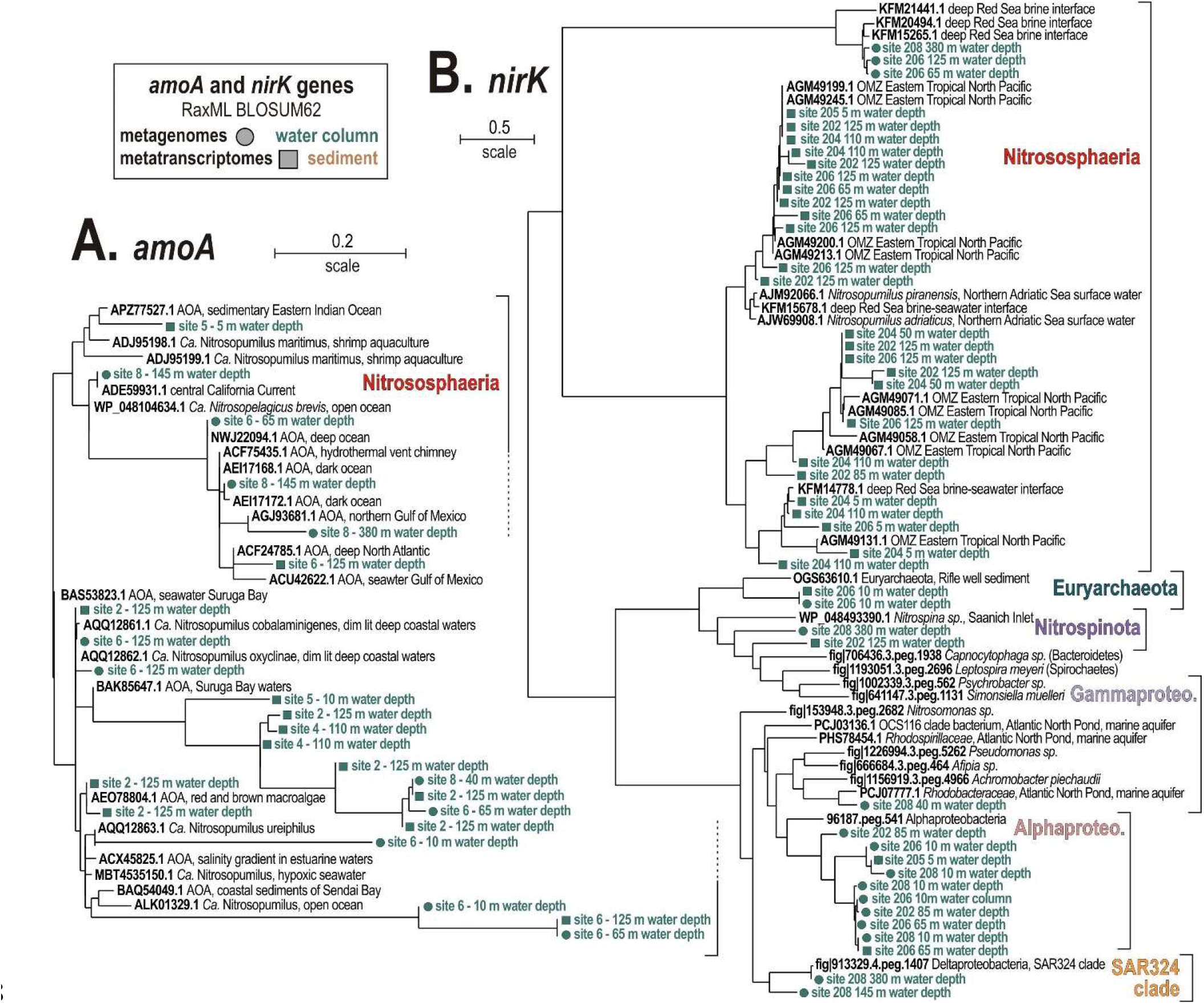
Phylogenetic analyses of predicted proteins encoded by (A) ammonia monooxygenase subunit A, and (B) nitrite reductase subunit K as the selected marker genes in the metagenomes and metatranscriptomes, based on RAxML using BLOSUM62 as the evolutionary model. (**A**) Phylogenetic tree of all *amoA* ORFs (185 aligned amino acid sites) detected in the metagenomes (circles) and metatranscriptomes (squares). (**B**) Phylogenetic tree of all *nirK* ORFs (133 aligned amino acid sites) detected in the metagenomes (circles) and metatranscriptomes (squares). Green and brown font respectively indicate water or sediment as the isolation source of the sequences.

### Carbon assimilation and sensitivity to oxygen depletion

Autotrophic and heterotrophic carbon assimilation in nitrifiers was assessed, on the one hand by looking for ORFs related to ribulose-1,5-diphosphate carboxylase (*RuBisCO*) in the Calvin-Benson-Bassham (CBB) cycle, ATP citrate lyase (*acly*) as the first step of the reductive tricarboxylic acid cycle (TCA) cycle, carbon monoxide dehydrogenase (*codh*) involved in CO oxidation and CO_2_ reduction, inclusive of the Wood-Ljungdahl (W-L) pathway, and acetyl-coenzyme A carboxylase (acc) as a step in the aerobic 3-hydroxypropionate/4-hydroxybutyrate (HP/HB) and anaerobic dicarboxylate/4-hydroxybutyrate (DC/HB) cycle as well as in anaplerotic CO_2_ assimilation (Erb, 2011); on the other hand by looking for acetyl-coenzyme A synthetase (*acs*) as the last metabolic step in glycolysis, and citrate synthase (cs) and pyruvate/lactate dehydrogenase (*pdh*) as the first metabolic steps in TCA cycle and lactate/pyruvate fermentation (Fig. 7).

**Figure 7.**
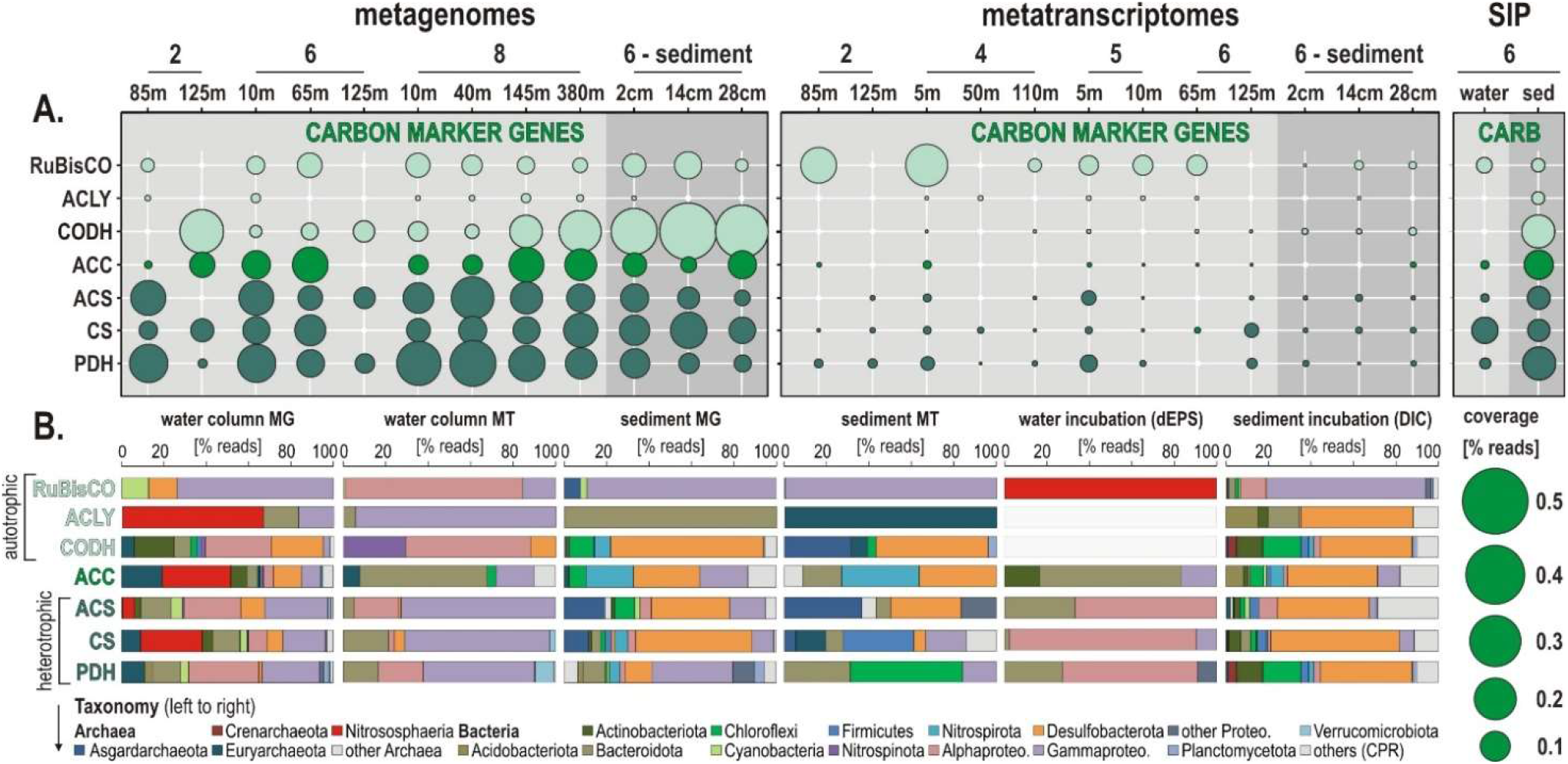
Metabolic functions and activities related to autotrophic and heterotrophic carbon assimilation in the water column, sediment and SIP incubations, and the corresponding taxonomic assignments at the phylum level. **(A)** Bubble plot showing the relative potential and expression of metabolic functions [% total reads] assigned to marker genes involved in autotrophic-heterotrophic carbon assimilation in the metagenomes, metatranscriptomes, stable isotope probing (SIP) water and sediment incubations (left to right). (**B**) Taxonomic bar charts [% reads] for the corresponding marker genes at the phylum level in the metagenomes (MG), metatranscriptomes (MT) and SIP incubations with water and ^13^C-labeled dEPS and with sediment and ^13^C-labeled bicarbonate (DIC). *Abbreviations:* RuBisCO: ribulose-1,5-diphosphate carboxylase (i.e. CBB cycle) / ACLY: ATP-citrate lyase (i.e. reductive TCA cycle) / CODH: carbon monoxide dehydrogenase (i.e. Wood-Ljungdahl pathway) / ACC: acetyl-coenzyme A carboxylase (i.e. HP/HB-DC/HP cycle) / ACS: acetylcoenzyme A synthetase (i.e. glycolysis) / CS: citrate synthase (e.g. TCA cycle) / PDH: pyruvate dehydrogenase (i.e. lactate/pyruvate fermentation).

ORF expression levels (i.e. % total transcripts) related to *RuBisCO* genes are highest in both oxic and dysoxic coastal waters (Fig. 7A) involving mostly the phyla Cyanobacteria and Protebacteria (Fig. 7B). In the SIP metagenomes incubated with water, *RuBisCO* genes are entirely assigned to Nitrososphaeria, whereas expression of *RuBisCO* genes in the seafloor and sediment SIP incubations is ruled by Gammaproteobacteria. In comparison, expression levels of ORFs assigned to *acly, codh* and *acc* genes are low, but tend to slightly increase offshore (sites 5 and 6). Heterotrophic processes assessed via transcription of *acs, cs* and *pdh* genes are detectable across all water column sampling sites and sediment, with generally higher levels of ORF expression in coastal waters than in the sediment (Fig. 7A). ORFs diagnostic of aerobic autotrophic carbon fixation (acc, i.e. HP/HB cycle) are expressed by pelagic taxa among Bacteroidota and Gammaproteobacteria, with little detection of Euryarchaeota and Chloroflexi, whereas in the sediment ORFs indicative of the anaerobic DC/HB cycle are expressed by Nitrospirota and Desulfobacterota.

In contrast, SIP metagenomes from water incubations only show ^13^C-labeling of the *acc* genes with taxonomic assignment to Bacteroidota. Assignments of ORFs related to glycolysis (acs) and TCA cycle (cs) indicate members of the Bacteroidota, Alpha- and Gammaproteobacteria as main actors of OM degradation in the water column and water incubations, and Euryarchaeota, Firmicutes and Desulfobacterota in the seafloor and sediment incubations (Fig. 7B). ORFs encoding genes for anaerobic fermentation (*pdh*), the W-L pathway (*codh*) and/or anaplerotic CO_2_ assimilation (Erb, 2011) are mostly expressed by Nitrospinota, Alpha- and Gammaproteobacteria in the OMZ, whereas in the sediment such OM fermentation processes are apparently driven by Chloroflexi and Bathyarchaeia.

ORFs assigned to genes encoding proteins for superoxide dismutase (*sod*), cytochromes cbb3, bd2, and b (*cyt b/d*) (Kalvelage et al., 2015), sensor histidine kinase (*shk*) and fumarate dehydrogenase (*frd*) were sorted to assess whether carbon assimilation pathways are associated with decreasing oxygen concentrations. Although metabolic potential based on % total reads tends to increase towards the oxycline, patterns of expression level remain inconspicuous, involving *shk* gene expression by Nitrososphaeria, Gammaproteobacteria and Bacteroidota in the water column (Supplementary Fig. 6).

## Discussion

### Active nitrifying and denitrifying populations along the Namibian coast

The taxonomic affiliation of 16S rRNA genes clearly separated two populations corresponding to the water column and sediment, the former being predominantly composed of diverse Proteobacteria, Cyanobacteria, Bacteroidota and Nitrososphaeria (Ulloa et al., 2013; Ganesh et al., 2014), whereas the latter consisted mainly of Chloroflexi, Nitrospirota and Desulfobacterota (Supplementary Fig. 5).

The relative abundances of presumed nitrogen-fixing populations were most abundant offshore. Those of nitrifying populations apparently thrived in OMZ waters on the shelf toward the north where lateral current increases ventilation of bottom waters (Ohde and Dadou, 2018), whereas denitrifying populations increased with water depth at all sites (Fig. 2). Consistent with this initial distribution of metabolic guilds (Fig. 2), the number of expressed ORFs encoding *amoA* and *nxr* genes involved in nitrification, and *nar*, *nir* and *nor* genes in denitrification increased from surface ocean down into OMZ waters (Fig. 4). Relative expression of NH_4_^+^, nitrite and urea transporters also increased down into OMZ waters along with nitroreductases and formate dehydrogenase (Einsle and Kroneck, 2004), which altogether points to concomitant metabolic activities in autotrophic, heterotrophic nitrification and denitrification (Fig. 4).

Based on phylogenetic analysis of 16S rRNA and *amoA* genes (Figs. 2A and 6A), pelagic AOA populations included taxa assigned to *Ca*. Nitrosopumilus (spp. maritimus, cobalaminigenes, oxyclinae, and ureiphilus) (Ijichi and Hamasaki, 2011) and *Ca*. Nitrosopelagicus (sp. brevis), whereas benthic ones were clearly distinct as they were all related to *Ca*. Nitrososphaera (sp. viennensis). Based on 16S rRNA genes, presumed AOB populations were affiliated with *Nitrosomonas* sp., *Nitrosococcus* sp. and *Nitrospira* sp. among respective phyla Gammaproteobacteria and Nitrospirota, in the absence of alphaproteobacterial sequences in the assemblage, e.g. *Nitrobacter* sp. (Fig. 3B). However, these bacteria were not identified in our phylogenetic analysis of *amoA* genes (Fig. 6A). Assignments of 16S rRNA genes also evidenced *Nitrospina-related* taxa as the prevalent pelagic NOB (Sun et al., 2019). Although both AOA and NOB were identified in taxonomic assemblages of the sediment, we could not detect any metabolic potential or activities towards benthic nitrification (Fig. 5).

Several sequences were affiliated with the anammox-related taxa Ca. Scalindua and Ca. Brocardia in the OMZ (Fig. 3C). However, these taxa were mostly present in the sediment and seemed not to be metabolically active as they could not be identified in the metatranscriptomes. At sites 2 and 3 where NH_4_^+^ may apparently diffuse out of the sediment, the water column profiles display intervals where NH_4_^+^ and nitrate are consumed while nitrate is being produced (Fig. 1B), which constitutes at least some geochemical suggestion that anammox processes may take place. At the SWI, Gammaproteobacteria prevailed in the taxonomic assemblage (Fig. 2), evidencing a benthic-pelagic transition where denitrification and DNRA can be coupled with sulfur oxidation (Callbeck et al., 2018) as sulfate reduction is the prevailing process in the underlying sediment (Vuillemin et al., 2022). Because nitrate is used as an oxidant by sulfur-oxidizing bacteria (Zhu et al., 2018) to drive DNRA, we suspect that anammox processes may be inhibited in the surface sediment (Jensen et al., 2008; Jin et al., 2013; Schunck et al., 2013). Instead, anammox-related Planctomycetota may be more metabolically active in mixed waters (Wasmund et al., 2016) when seasonal OM production and particle sinking are highest (Karthäuser et al., 2021) and nitrite is replenished (Brüchert et al., 2003; Callbeck et al., 2021).

Altogether, variations in the related ORF expression patterns (Fig. 3) suggest that trophic interactions in austral winter are more determined by aerobic remineralization of sinking OM to NH_4_^+^ (Füssel et al., 2012; Kalvelage et al., 2015) on the shelf with limited benthic releases in the benthic boundary layer. The fact that NH_4_^+^ concentrations systematically decline in the OMZ at oxygen concentrations <60 μM (Figs. 1B–1C) argues for nitrifier denitrification processes in the OMZ upper waters sustained by lateral transport along the Namibian coast (Inthorn et al., 2006b, 2006a).

### Nitrifier denitrification is the prevailing process in dysoxic waters

In general, our profiling of expressed ORFs confirmed the deficit in nitrogen fixation in the sunlit ocean (Ulloa et al., 2013) as the related genes (*nif*) were minor and restricted to surface waters (Fig. 4) even though it involved a taxonomically diverse pelagic community (Gier et al., 2016; Jayakumar and Ward, 2020), e.g. Cyanobacteria, Alpha-, Gammaproteobacteria, Firmicutes, Euryarchaeota (Fig. 5B). ORFs actively encoding aerobic *amo* and *nxr* genes consistently increased from the surface ocean down into the OMZ at all sites (Fig. 4), displaying high expression levels concomitant with NH_4_^+^ transporters (Fig. 5A). Metabolic use of other forms of fixed nitrogen through enzymes like *nmo* genes and ureases was only detectable in surface ocean of the shelf, which confirmed that ureaderived ammonia oxidation contributes little to nitrification activities in the coastal OMZ in comparison to NH_4_^+^-limited waters offshore (Tolar et al., 2017; Kitzinger et al., 2019; Shiozaki et al., 2021). Genes related to nitrilases and cyanases were identified, but not expressed (Fig. 4). ORFs encoding genes for nitrification were not detected in the sediment (Fig. 4). These patterns show that the upper layer of OMZ waters is critical for microaerobic nitrifying microorganisms, particularly the AOA and NOB (Kitzinger et al., 2020).

*Ca*. Nitrosopumilus and *Ca*. Nitrosopelagicus (Figs. 3A and 6A) have a demonstrated capacity for growth using ammonia oxidation as an energy source (Vuillemin et al., 2019), resulting in stoichiometric production of nitrite (Könneke et al., 2005; Li et al., 2013) that can be re-oxidized to nitrate (Lam and Kuypers, 2011; Füssel et al., 2012). A relatively high number of ORFs was assigned to *nxr* genes, whose taxonomic assignments (Fig. 5B) showed that Nitrospinota is the only phylum actively involved in processes of nitrite oxidation to nitrate (Rani et al., 2017). This shows that ammonia was actively oxidized to nitrite by archaea and to nitrate by bacteria (Figs. 4 and 5A), leading to reciprocal feeding interactions in the vicinity of the OMZ (Fig. 1C), which may limit the loss of fixed nitrogen via denitrification or anammox processes (Fernandez and Farias, 2012; Koch et al., 2015; Mueller et al., 2021). Consistent with the presence of *Nitrospira* (sp. *moscoviensis*) in 16S rRNA genes but absence of any related bacterial *amoA* gene, we did not identify ORFs encoding hydroxylamine dehydrogenase (i.e. *hao* cluster), implying that complete nitrification did not proceed via comammox (Daims et al., 2016; Palomo et al., 2018; Long et al., 2021). Instead, the produced nitrite appeared to be actively reduced via transcription of *nirK* genes (Braker Gesche et al., 2000) by the aforementioned Nitrososphaeria, and Nitrospinota, as well as Desulfobacterota in the OMZ (Fig. 4A–4B). ORFs assigned to anaerobic *nir* genes had by far the highest expression level in terms of nitrogen reduction compared to *nar* genes (Fig. 5A), which demonstrates that nitrifier denitrification was the prevailing process from the surface ocean into OMZ waters across all sites (Figs. 4 and 5) in spite of nitrate-replete OMZ waters (Fig. 1C). The taxonomic assemblage of the metabolic guild actively expressing *nar* and *nir* genes (Smith et al., 2007, 2015) in the water column included Nitrospinota, Nitrososphaeria, Gammaproteobacteria and fewer Desulfobacterota (Fig. 5B). Some ORFs assigned to *nor* genes were subsequently expressed in deep waters (Fig. 4), mostly by Gammaproteobacteria, Desulfobacterota and Nitrospirota (Fig. 5B) albeit at much lower levels. Interestingly, *nos* genes which represents the final step of denitrification (i.e. N_2_O reduction to N_2_) were only detectable in the surface ocean where they were assigned to Bacteroidota and fewer Nitrospirota and Proteobacteria (Fig. 5B) known to display atypical N_2_O-scavenging abilities (Wyman et al., 2013; Bertagnolli et al., 2020).

These expression patterns show that nitrifiers were most active in transcribing *nirK* genes (Lawton et al., 2013), potentially leading to N_2_O production (Stein, 2011b; Ji et al., 2015; Buitenhuis et al., 2018; Han et al., 2022) due to the necessity to detoxify the produced nitric oxide in the OMZ rather than through canonical denitrification (Stein, 2011a; Bertagnolli et al., 2020). In addition, nitroreductases were expressed in dysoxic waters, but only locally (site 4). Thus, in spite of active recycling of metabolites (urea, R-NO_2_, NH_4_^+^) via microaerobic microbial respiration (Behrendt et al., 2013; Kitzinger et al., 2019), our results support the general deficit of fixed nitrogen as an electron acceptor (Kraft et al., 2011) and nitrifier denitrification (Fernandez and Farias, 2012; Lawton et al., 2013) as the main nitrogen reduction pathway in the water column at ca. 60 μM oxygen concentrations (Fig. 1), potentially leading to N_2_ and N_2_O gas emissions (Wrage-Mönnig et al., 2018).

### Ammonification and denitrification in sulfidic sediment

When the SWI of the Namibian inner shelf is anoxic, H_2_S, CH_4_ and NH_4_^+^ frequently diffuse out of the sediment forming sulfur plumes in the water column during austral summer (Brüchert et al., 2003, 2009; Neumann et al., 2016). As stratification decreases along the shelf, NH_4_^+^ can be locally released from the seafloor into the benthic boundary layer (Füssel et al., 2012; Neumann et al., 2016). Seasonal variations in Namibian coastal waters may thereby promote chemolithoautotrophy with sulfur and NH_4_^+^ as successive electron donors (Farías et al., 2009; Lam and Kuypers, 2011) to drive sulfur-dependent denitrification by Gammaproteobacteria (Lipsewers et al., 2017; Vuillemin et al., 2022) pursued by nitrifier denitrification by Nitrososphaeria and Nitrospinota in the water column (Wrage-Mönnig et al., 2018; Ruiz-Fernández et al., 2020).

Taxa identified as presumed nitrifiers in the sediments were closely affiliated with *Ca*. Nitrososphaera viennensis (Fig. 3A), *Nitrosomonas nitrosa*, and candidate clades among *Nitrosococcaceae* and *Nitrospinaceae* (Fig. 3B). However, metabolic activities towards oxidation of nitrogen compounds only involved *nmo* genes by *Lokiarchaeum* sp. (Fig. 5B), potentially resulting in minor production of nitrite (Zhang et al., 2022). Consistent with the apparent lack of metabolic expression by anammox-related Planctomycetota (Fig. 3C), we did not detect any ORF related to hydrazine (i.e. H_2_N_4_) oxidoreductase (hzo) in shallow sediment (Kong et al., 2013). In spite of members of the Gammaproteobacteria, e.g. *Beggiatoa, Thiothrix, Thioploca* (Vuillemin et al., 2022), being capable of coupling DNRA with sulfide oxidation (Schunck et al., 2013; Crowe et al., 2018; Zhu et al., 2018), the abundance of ORFs expressing ammonia-forming *nrf* genes (i.e. DNRA) was minor (Figs. 4 and 5), solely involving taxa among Nitrospirota in sulfidic sediments (Murphy et al., 2020).

In highly reducing nitrate-depleted sediment (Supplementary Fig. 4), the survival of denitrifiers is generally poor, and such populations are likely to employ Fe^3+^ reduction in their energy metabolism (Coby Aaron J. et al., 2011), which is theoretically absent in Namibian sediment (Böning et al., 2020), thus arguing for another pathway of NH_4_^+^ production. Expression of nitroreductases by Bacteroidota suggests active degradation of OM with release of nitro (i.e. R-NO_2_) compounds (Roldán et al., 2008). We deduct that benthic NH_4_^+^ production (Fig. 1D) potentially released into the coastal OMZ waters (Fig. 1C) results from amino acid remineralization (i.e. algal necromass) via expression of proteases by heterotrophic bacteria (Neumann et al., 2016) rather than DNRA (Pantoja and Lee, 2003; Vuillemin et al., 2022). In the bottom part of the core (28 cmbsf), ORFs assigned to *nar* and *nir* genes were mainly expressed by Proteobacteria, whereas *nor* and *nos* genes were also expressed by taxa among Methylomirabilota (former candidate NC10) (Padilla et al., 2016) and Desulfobacterota, but gene expression of these final steps of denitrification was minor in the sediment (Figs. 4 and 5B).

Results from water incubations with dEPS over an 18-hours period confirmed remineralization of algal necromass as all the genes involved in the initial reduction of fixed nitrogen (i.e. *nif, nar, nir*, NH_4_^+^ transporters) were ^13^C-labeled in the total absence of ammonia oxidation (Fig. 4). Bacteroidota appeared as the main aerobic degraders of algal necromass that rapidly grew and led to microoxic conditions under which processes of denitrification initiated, as shown by ^13^C-labeling of *nif, nir*, and *ntr* genes (Fig. 5C). Other taxa that were ^13^C-labeled for nitrate, NH_4_^+^ and urea transporters were found among Planctomycetota and Proteobacteria, respectively (Fig. 5C). Sediment incubations with DIC over a 10-days period resulted in the ^13^C-labeling of all the genes involved in denitrification and DNRA (Figs. 4 and 5A). The related nitrate-reducing taxa included a majority of Desulfobacterota and Gammaproteobacteria, with fewer Chloroflexi, Nitrospinota and Actinobacterota (Fig. 5B). Under *in vitro* conditions, the highest number of ^13^C-labeled ORFs were assigned to proteases (Vuillemin et al., 2022), indicating that organic nitrogen was assimilated from algal necromass via fermentative processes.

### Autotrophic or heterotrophic nitrifier denitrification?

Under oxic conditions, AOA and NOB have metabolic potential for autotrophic carbon fixation through the HP/HB and CBB cycle (Könneke et al., 2014; Alfreider et al., 2018). In spite of being common aerobes, Nitrospinota have in theory the capability to fix carbon via the reductive TCA (Luecker et al., 2013). Expressed ORFs encoding the *RuBisCO, acly* and *acc* genes in the water column were mostly assigned to Alpha- and Gammaproteobacteria and Bacteroidota (Fig. 7). Autotrophic carbon fixation by Nitrosophaeria was either not expressed or below detection at the present sequencing depths, whereas nitrite-oxidizing Nitrospinota could apparently evolve as potential autotrophs by expressing *codh* genes (Zhang et al., 2020) instead of the reductive TCA as previously reported (Luecker et al., 2013). Interestingly, results from SIP metagenomes incubated with water and dEPS revealed ^13^C-labeling of *RuBisCO* genes that were exclusively assigned to Nitrosophaeria (Fig. 7B). Along with our profiling of ORF expression, this confirms that chemoautotrophic ammonia and nitrite oxidizers can adapt to low concentrations of fixed nitrogen in heterotrophic food web by coupling nitrogen and carbon cycling using simple organic substrates (Kitzinger et al., 2019, 2020), e.g. urea (CH_4_N_2_O), cyanate (CH_3_OCN), amino acids and nitro compounds (R-NO_2_), for heterotrophic nitrification and nitrifier denitrification (Stein, 2011b; Dekas et al., 2019), potentially with anaplerotic CO_2_ assimilation mediated by *RuBisCO* and *codh* genes (Pester et al., 2011; Koch et al., 2015; Aylward and Santoro, 2020; Reji and Francis, 2020; Zhang et al., 2020). Consistent with oxygen drawdown during heterotrophic degradation of algal OM sinking in the water column, expressed ORFs related to *acs*, *cs* and *pdh* genes indicated the prevalence of fermentative glycolysis and TCA cycle for carbon assimilation under dysoxic conditions (Fig. 7). In this context, ORFs assigned to formate dehydrogenase, which is also the first step of the respiratory chain in denitrification (Einsle and Kroneck, 2004), were increasingly expressed down into the OMZ by Alpha- and Gammaproteobacteria (Figs. 4 and 5). Although not presently expressed by nitrifiers, the electron transfer from formate to nitrate may support versatility in the use of simple carbon resources during denitrification as formate can be readily oxidized through nitrate reduction for anaerobic growth under anoxia (Koch et al., 2015; Daims et al., 2016).

## Conclusions

Altogether metatranscriptomic profiling demonstrates that denitrification clearly outcompassed nitrogen fixation and ammonia-forming nitrate reduction, or DNRA. Nitrifiers, such as ammonia-oxidizing Nitrososphaeria and nitrite-oxidizing Nitrospinota, displayed high expression levels of *amo* and *nxr* genes in dysoxic waters in concomitance with *nirK* genes, thereby also performing nitrite reduction. These two groups of nitrifiers appeared to accessorize a certain degree of mixotrophy under OMZ conditions by microaerobically oxidizing simple organic compounds and anaplerotic CO_2_ assimilation coupled with nitrite reduction. Intriguingly, ORFs related either to anammox or comammox were not detected. Most likely, the nitrite necessary to anammox and comammox reactions was either re-oxidized to nitrate or reduced to nitric oxide by Nitrososphaeria and Nitrospinota during nitrifier denitrification. The subsequent reduction from nitric to nitrous oxide was driven by Nitrospirota and Gammaproteobacteria in OMZ waters, whereas the produced N_2_O was scavenged at the ocean surface. To conclude, in austral winter the main pathways to potential N_2_O production stem from OM remineralization by a microbial consortium actively performing heterotrophic nitrification and nitrifier denitrification in the vicinity of the oxycline.

## Data Availability

Geochemical data (Siccha and Kucera, 2018; Ferdelman et al., 2021b, 2021a; Garaba et al., 2021) are archived and publicly available from the PANGAEA^®^ Data Publisher for Earth and Environmental Science (dataset #895640, #931090, #931097, and #928943). Bathymetric data (Weatherall et al., 2021) are publicly available (https://www.bodc.ac.uk/data/published_data_library) from the GEBCO_2022 Grid.

All scripts and codes used to produce sequence analyses have been posted on GitHub with a link to the instructions on how to conduct the scripts (github.com/williamorsi/MetaProt-database). All metagenome, metatranscriptome and 16S rRNA gene data are publicly accessible in NCBI through BioProject number PRJNA525353.

## Author Contributions

AV conceived the idea for the study, extracted nucleic acids, performed quantitative PCR assays, library preparation and Illumina sequencing, analyzed the data, designed the figures and wrote the paper.

## Acknowledgments

W.D. Orsi and Ö.K. Coskun are acknowledged for their help with bioinformatics and for providing feedback on the manuscript. The crew of the F/S Meteor and organizing committee of the oceanographic expedition are acknowledged in obtaining samples and producing geochemical data.

