## Supplementary for "Nitrogen cycling activities during decreased stratification in the coastal oxygen minimum zone off Namibia"

The present supplemental file is part of a non-peer reviewed preprint submitted to **bioRxiv.org**. Please note that the related manuscript will be submitted to a scientific journal for peer review. Subsequent versions of this supplemental file may have slightly different content. If accepted, the final version of the manuscript and supplemental file will be available via the "Peer-reviewed Publication DOI" link on the right-hand side of this webpage. Please feel free to contact the corresponding author, your feedback is welcome.

### ***Supplementary Material***

**Content:**


- **Supplementary Figure 1.** Pictures of sediment core
- **Supplementary Figure 2.** $^{13}C$-labeled taxonomic assemblages and qPCR buoyant density curves
- **Supplementary Figure 3.** Additional geochemical profiles for the water column at site 2, 3, 4 and 6
- **Supplementary Figure 4.** Pore water geochemical profiles and 16S rRNA genes in sediments
- **Supplementary Figure 5.** Taxonomy of 16S rRNA genes, metagenomes, and transcriptomes
- **Supplementary Figure 6.** Metabolic functions and activities related to putative oxygen sensors

- **Supplementary Table 1.** Sequencing and assembly statistics of metagenomes and transcriptomes
- **Supplementary Table 2.** Binning statistics for metatranscriptomes

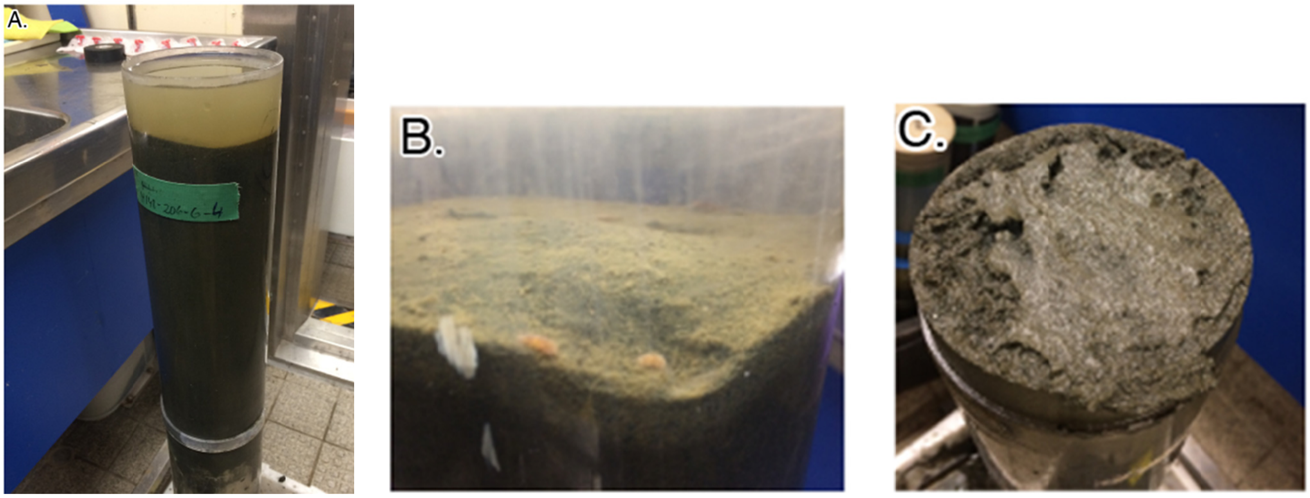


**Supplementary Figure 1. Sediment cores retrieved from the Namibian shelf.** (**A**) Pristine core recovered, together with bottom water. (**B**) Zoom in on the core top surface evidencing an undisturbed sediment-water interface. (**C**) Photo of core inside at ca. 15 cm depth, taken during sectioning. The sediment is composed primarily of green mud and foraminiferal sand. Photos are reproduced from (Orsi et al., 2020b, 2020a).

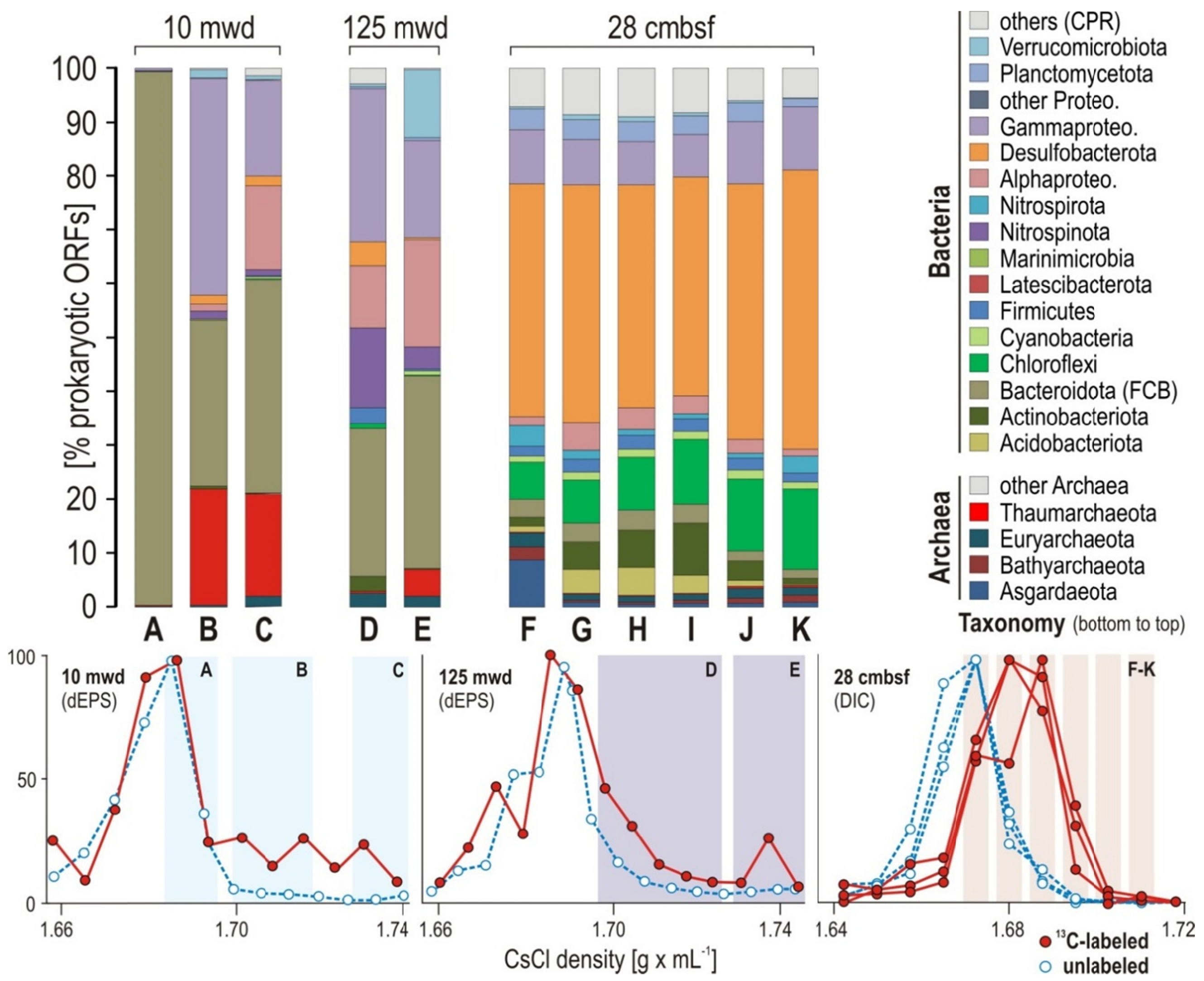


**Supplementary Figure 2. Taxonomic assemblages from the total $^{13}$C-labeled "isotopically heavy" DNA fractions identified via qPCR buoyant density curves.** (**Top**) Taxonomic assemblages of the "heavy" DNA fractions [% prokaryotic ORFs] considered indicative of phyla showing $^{13}$C-dEPS and $^{13}$C-bicarbonate assimilation. (**Bottom**) qPCR buoyant density curves for 16S rRNA genes obtained from DNA fractions of the 18 hours SIP incubations with water from 10 and 125 mwd and $^{13}$C-dEPS, and 10 days SIP incubations with sediment from 28 cmblf at site 6 and $^{13}$C-bicarbonate (red dots: $^{13}$C-labeled; blue dots: unlabeled controls). Illustration is modified after (Vuillemin et al., 2022).

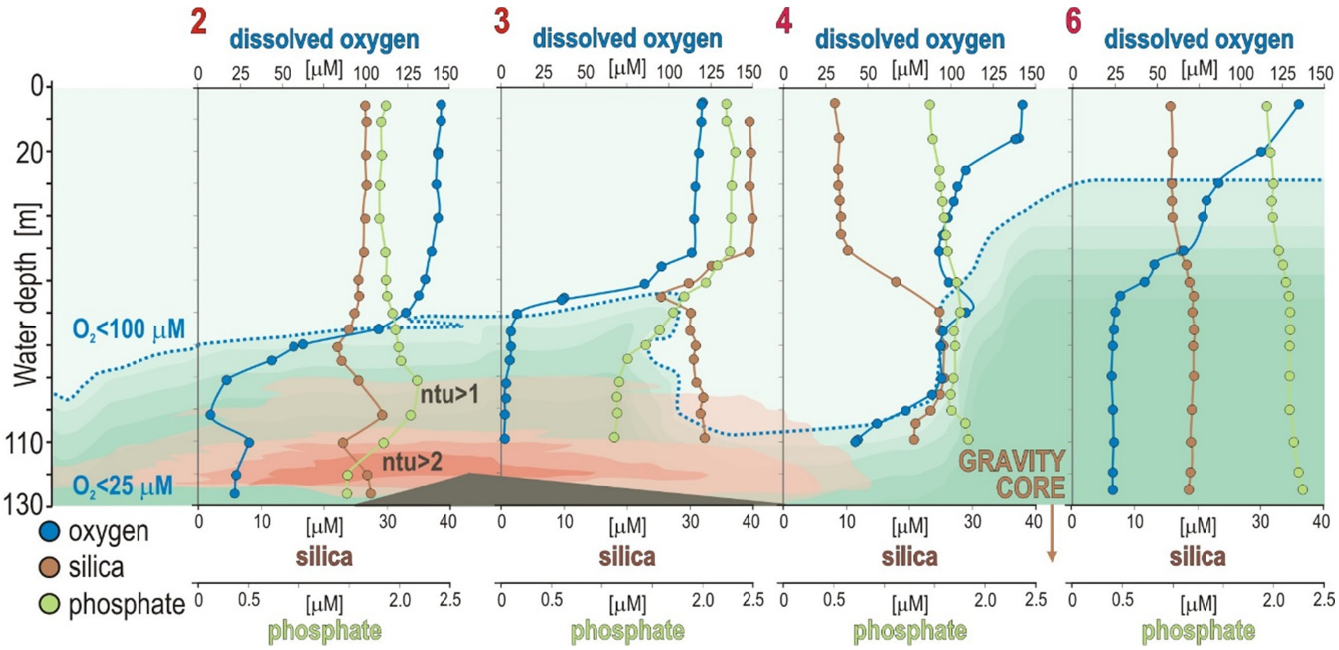


**Supplementary Figure 3. Geochemical profiles for the water column at site 2, 3, 4 and 6.** Concentration profiles for dissolved oxygen (blue) with an oxycline defined as <100 μM (dotted line), silica (brown), phosphate (green) and turbidity (pink) in nephelometric turbidity units (ntu) at each successive sampling site. Water column data are from (Siccha and Kucera, 2018; Ferdelman et al., 2021b).

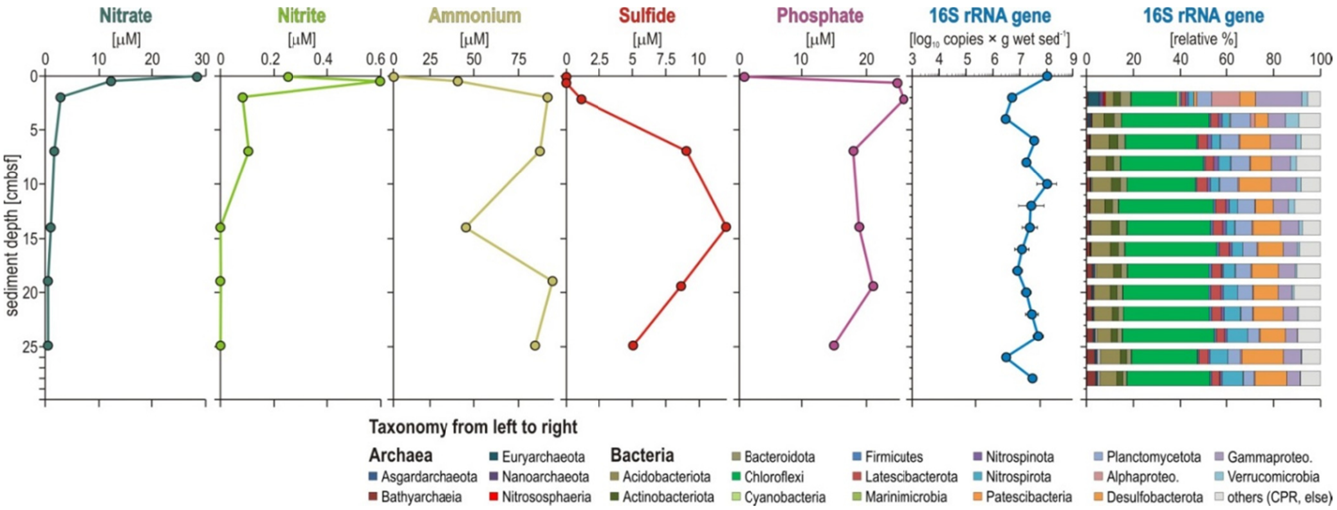


**Supplementary Figure 4. Geochemical profiles for the sediment at site 6 with 16S rRNA gene density and diversity.** From left to right: Geochemical profiles of nitrate, nitrite, ammonium, sulfide and phosphate [µM], followed by 16S rRNA gene density [$log_{10}$ gene copies × g wet $sed^{-1}$] assessed via quantitative PCR and 16S rRNA gene taxonomic diversity [relative %]. Pore water data are from (Ferdelman et al., 2021a).

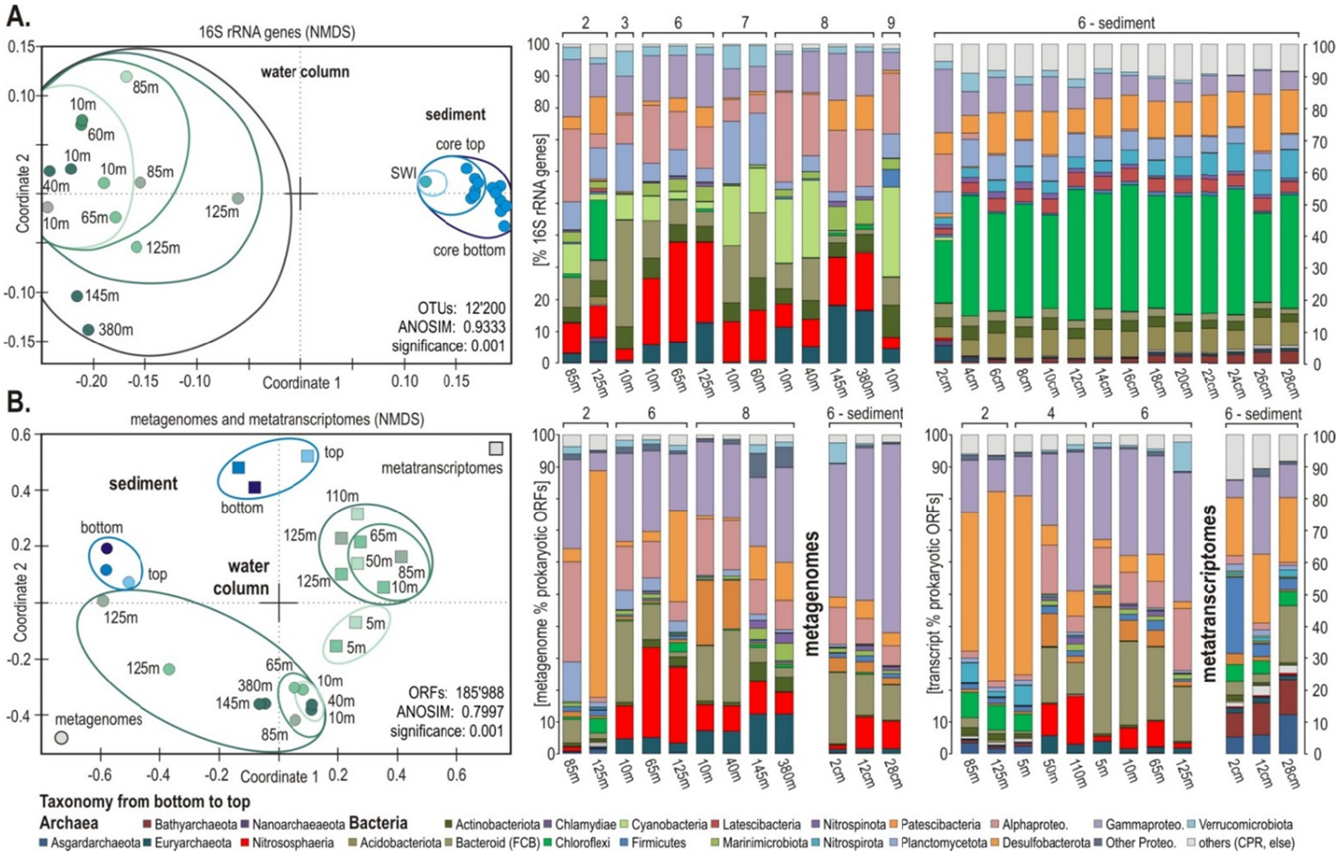


**Supplementary Figure 5. Beta diversity and taxonomic assemblages of 16S rRNA gene amplicons, metagenomes and metatransciptomes.** (**A**) Non-metric multidimensional scaling (NMDS) plot based on all OTUs across all water column and sediment samples and their corresponding taxonomic assemblages [% 16S rRNA genes]. **(B**) NMDS plot based on all prokaryotic ORFs from the metagenomes (circles) and metatranscriptomes (squares) and their corresponding taxonomic assignments [% ORFs]. Illustration is modified from (Vuillemin et al., 2022).

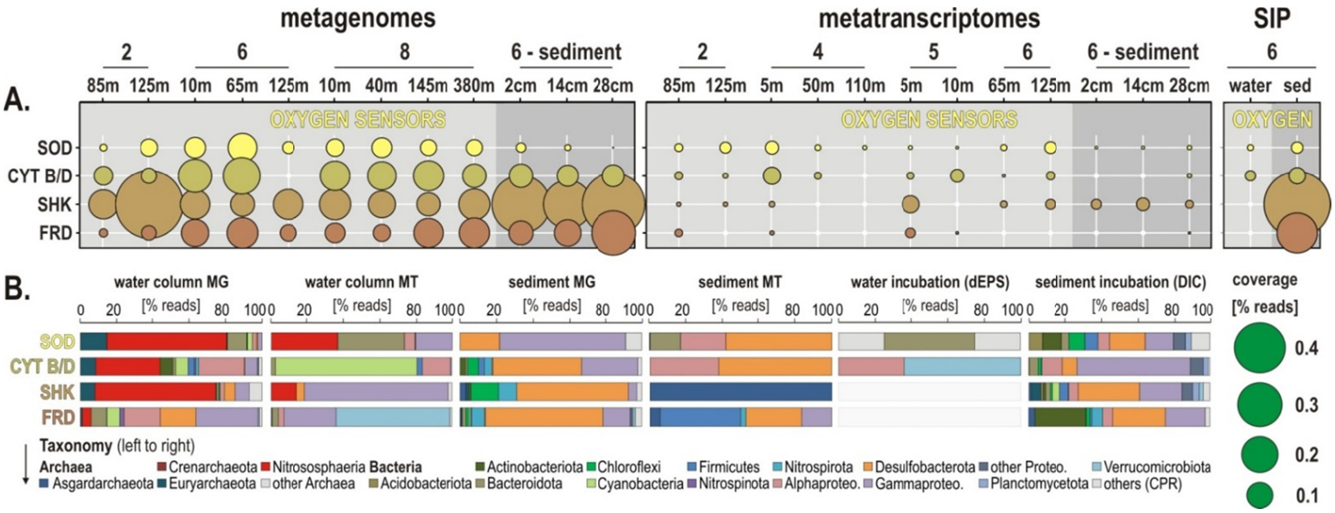


**Supplementary Figure 6. Metabolic functions and activities related to putative oxygen sensors in the water column, sediment and SIP incubations, and the corresponding taxonomic assignments at the phylum level. (A)** Bubble plot showing the relative potential and expression level of metabolic functions [% total reads] assigned to marker genes encoding putative oxygen sensors in the metagenomes, metatranscriptomes, stable-isotope probing (SIP) water and sediment incubations (left to right). (**B**) Taxonomic bar charts [% reads] for the corresponding marker genes at the phylum level in the metagenomes (MG), metatranscriptomes (MT) and SIP incubations with water and $^{13}C$-labeled dEPS and with sediment and $^{13}C$-labeled bicarbonate (DIC). *Abbreviations:* SOD: superoxide dismutase / CYT B/D: cytochromes cbb3, bd2 and b / SHK: sensor histidine kinase / FRD: fumarate reductase.

**Supplementary Table 1. Sequencing and assembly statistics for metatranscriptomes, metagenomes, and SIP-metagenomes.**

METATRANSCRIPTOMES

| Station | Sample type | oxygen level | Depth (m) | Replicate | # reads (millions) | # contigs | reads mapping (millions) | % of reads mapping to contigs |
|---|---|---|---|---|---|---|---|---|
| site-202 | water column | oxycline | 85 | a | 3 | 13,274 | 2.1 | 70.0 |
| | | | | b | 3.6 | 27,862 | 2.2 | 61.1 |
| | | OMZ | 125 | a | 4.7 | 19,254 | 3.5 | 74.5 |
| | | | | b | 3.4 | 15,817 | 2.5 | 73.5 |
| site-204 | water column | Surface (oxic) | 5 | a | 10.2 | 67,010 | 8.2 | 80.4 |
| | | | | b | 9.1 | 41,408 | 7 | 76.9 |
| | | oxycline | 50 | a | 9.4 | 21,960 | 8.2 | 87.2 |
| | | | | b | 7.6 | 17,647 | 6.6 | 86.8 |
| | | OMZ | 110 | a | 13.8 | 27,168 | 12.2 | 88.4 |
| | | | | b | 7.9 | 29,864 | 6.2 | 78.5 |
| site-206 | water column | oxic | 5 | a | 5.7 | 19,334 | 4.6 | 80.7 |
| | | | | b | 6.7 | 37,547 | 5.1 | 76.1 |
| | | | 10 | a | 7.2 | 20,735 | 6.3 | 87.5 |
| | | | | b | 3.7 | 10,464 | 3.2 | 86.5 |
| | | oxycline | 65 | a | 7.2 | 16,298 | 6.1 | 84.7 |
| | | | | b | 3.5 | 9,860 | 2.7 | 77.1 |
| | | OMZ | 125 | a | 1.7 | 5,526 | 1.3 | 76.5 |
| | | | | b | 5.3 | 16,420 | 4.2 | 79.2 |
| | | | | c | 7.7 | 25,788 | 6.1 | 79.2 |
| | sediments | hypoxic | core top | a | 4.6 | 2,602 | 3.7 | 80.4 |
| | | | | b | 11.1 | 2,927 | 9.2 | 82.9 |
| | | sulfidic | 12 cm | a | 3.8 | 4,362 | 2.7 | 71.1 |
| | | | | b | 2.2 | 2,726 | 1.3 | 59.1 |
| | | | | c | 3.4 | 5,888 | 2.1 | 61.8 |
| | | | 28 cm | a | 3.8 | 7,429 | 2.3 | 60.5 |
| | | | | b | 5.7 | 9,636 | 4.2 | 73.7 |
| | | | | c | 4.1 | 5,660 | 2.7 | 65.9 |

METAGENOMES

| Station | Sample type | oxygen level | Depth (m) | Replicate | # reads (millions) | # contigs | reads mapping (millions) | % of reads mapping to contigs |
|---|---|---|---|---|---|---|---|---|
| site-202 | water column | oxycline | 85 | a | 5.5 | 79,040 | 1.7 | 30.9 |
| | | OMZ | 125 | a | 3.6 | 81,412 | 0.6 | 16.7 |
| site-206 | water column | oxic | 10 | a | 4.9 | 102,742 | 1.5 | 30.6 |
| | | oxycline | 65 | a | 5.8 | 95,018 | 1.2 | 20.7 |
| | | OMZ | 125 | a | 7.1 | 49,885 | 0.88 | 12.4 |
| | sediments | hypoxic | core top | a | 15.7 | 80,933 | 1.26 | 8.0 |
| | | sulfidic | 12 cm | a | 5.1 | 12,580 | 0.15 | 2.9 |
| | | | 28 cm | a | 3.6 | 9,583 | 0.12 | 3.3 |
| site-208 | water column | oxic | 10 | a | 5 | 70,549 | 1.2 | 24.0 |
| | | | 145 | a | 7.8 | 125,658 | 1.9 | 24.4 |
| | | | 380 | a | 5 | 79,761 | 1.4 | 28.0 |

SIP-metagenomes

| Station | SIP incubation | Sample type | oxygen level | Depth (m) | $^{13}$C-SIP metagenome | Density range (g mL$^{-1}$) | # reads (millions) |
|---|---|---|---|---|---|---|---|
| site-206 | $^{13}$C-dEPS | water column | oxic | 10m | 10m_a | 1.685-1.70 | 4.9 |
| | | | | | 10m_b | 1.70-1.74 | 4.7 |
| | | | | | 10m_c | 1.74-1.76 | 3.7 |
| | | | OMZ | 125m | 125m_a | 1.70-1.73 | 5.8 |
| | | | | | 125m_b | 1.73-1.745 | 2.7 |
| | $^{13}$C-bicarbonate | sediment | sediment | 23 cm | 23cm_a | 1.682 | 9.6 |
| | | | | | 23cm_b | 1.689 | 2.7 |
| | | | | | 23cm_c | 1.697 | 15.5 |
| | | | | | 23cm_d | 1.705 | 12.8 |
| | | | | | 23cm_e | 1.713 | 8.7 |
| | | | | | 23cm_f | 1.722 | 1.6 |

**Supplementary Table S2. Binning statistics for metatranscriptomes.**

| Sample | total_num_reads | num_INDELs_reported | total_reads_kept | num_SNVs_reported | total_reads_mapped | percent_mapped |
|---|---|---|---|---|---|---|
| Site 202 85m (MT) | 5994376 | 7390 | 2543311 | 163023 | 2543311 | 42.43 |
| Site 202 125m (MT) | 6646454 | 10235 | 3257803 | 245540 | 3257803 | 49.02 |
| Site 204 5m (MT) | 17164574 | 12126 | 8786590 | 259866 | 8786590 | 51.19 |
| Site 204 50m (MT) | 15809144 | 13804 | 8992028 | 321241 | 8992028 | 56.88 |
| Site 204 110m (MT) | 21100648 | 9460 | 15912601 | 161755 | 15912601 | 75.41 |
| Site 206 5m (MT) | 11629756 | 8151 | 5088200 | 139498 | 5088200 | 43.75 |
| Site 206 10m (MT) | 11399926 | 14046 | 5529901 | 320131 | 5529901 | 48.51 |
| Site 206 65m (MT) | 9887270 | 11220 | 5402852 | 249057 | 5402852 | 54.64 |
| Site 206 125m (MT) | 20230136 | 13643 | 10912948 | 344857 | 10912948 | 53.94 |
| Site 206 core top3 (MT) | 3393648 | 7991 | 1706512 | 111746 | 1706512 | 50.29 |
| Site 206 core top (MT) | 9222134 | 12171 | 4416491 | 172450 | 4416491 | 47.89 |
| Site 206 core 12cm (MT) | 8446572 | 9040 | 2695838 | 213050 | 2695838 | 31.92 |
| Site 206 28cm (MT) | 59101508 | 17092 | 6830510 | 943736 | 6830510 | 11.56 |

| bins | total_length | num_contigs | N50 | GC_content | percent_completion | percent_redundancy | t_domain | t_phylum | t_class | t_order | t_family | t_genus | t_species |
|---|---|---|---|---|---|---|---|---|---|---|---|---|---|
| MAXBIN__031 | 6449290 | 3961 | 1549 | 52.0 | 91.55 | 67.61 | Bacteria | Desulfobacterota | Syntrophobacteria | BM002 | BM002 | BM002 | BM002 sp002899795 |
| MAXBIN__034 | 6583881 | 4089 | 1513 | 63.6 | 87.32 | 88.73 | Bacteria | Actinobacteriota | Acidimicrobiia | UBA5794 | UBA4744 | UBA4744 | UBA4744 sp002403855 |
| MAXBIN__036 | 3542706 | 2068 | 1711 | 64.1 | 67.61 | 32.39 | Bacteria | Myxococcota | | | | | |
| MAXBIN__033 | 2915624 | 1980 | 1428 | 61.7 | 63.38 | 35.21 | Bacteria | Myxococcota | Polyangia | Polyangiales | SG8-38 | SG8-38 | SG8-38 sp003647035 |
| MAXBIN__019 | 738826 | 497 | 1426 | 47.0 | 60.56 | 92.96 | Bacteria | | | | | | |
| MAXBIN__003 | 2410097 | 1526 | 1558 | 45.3 | 54.93 | 21.13 | Bacteria | Proteobacteria | Gammaproteobacteria | Enterobacterales | Alteromonadaceae | Pseudoalteromonas | |
| MAXBIN__009 | 434046 | 258 | 1616 | 41.1 | 50.70 | 53.52 | Bacteria | Proteobacteria | Gammaproteobacteria | PS1 | Thioglobaceae | Thioglobus | Thioglobus singularis |
| MAXBIN__018 | 102837 | 66 | 1513 | 32.6 | 50.70 | 36.62 | Bacteria | Bacteroidota | Bacteroidia | Flavobacteriales | Flavobacteriaceae | MAG-121220-bin8 | MAG-121220-bin8 sp004214185 |
| MAXBIN__032 | 2237305 | 1428 | 1537 | 46.9 | 47.89 | 11.27 | Bacteria | Desulfobacterota | Syntrophobacteria | BM002 | BM002 | BM002 | BM002 sp002899795 |
| MAXBIN__025 | 489338 | 327 | 1400 | 30.7 | 47.89 | 43.66 | Bacteria | Bacteroidota | Bacteroidia | Flavobacteriales | | | |
| MAXBIN__035 | 2493781 | 1879 | 1276 | 69.4 | 42.25 | 19.72 | Bacteria | Myxococcota | UBA9160 | UBA9160 | UBA6930 | UBA6930 | UBA6930 sp002450755 |
| MAXBIN__024 | 233551 | 157 | 1386 | 43.5 | 42.25 | 23.94 | Bacteria | | | | | | |
| MAXBIN__007 | 267590 | 156 | 1683 | 32.2 | 39.44 | 4.23 | | | | | | | |
| MAXBIN__021 | 1617878 | 1099 | 1373 | 39.7 | 38.03 | 35.21 | Bacteria | Proteobacteria | Gammaproteobacteria | PS1 | Thioglobaceae | Thioglobus | Thioglobus sp001628405 |
| MAXBIN__026 | 1308179 | 908 | 1368 | 42.4 | 36.62 | 22.54 | | | | | | | |
| MAXBIN__011 | 106535 | 48 | 2371 | 45.0 | 32.39 | 4.23 | Bacteria | Proteobacteria | Gammaproteobacteria | Pseudomonadales | Porticoccaceae | HTCC2207 | |
| MAXBIN__013 | 944322 | 704 | 1286 | 49.2 | 30.99 | 11.27 | Bacteria | Proteobacteria | Gammaproteobacteria | Pseudomonadales | | | |
| MAXBIN__016 | 222421 | 166 | 1293 | 37.2 | 29.58 | 22.54 | Bacteria | SAR324 | SAR324 | SAR324 | NAC60-12 | Arctic96AD-7 | Arctic96AD-7 sp002685535 |
| MAXBIN__022 | 133005 | 86 | 1570 | 38.4 | 28.17 | 19.72 | Bacteria | | | | | | |
| MAXBIN__014 | 177509 | 132 | 1304 | 32.9 | 26.76 | 28.17 | Bacteria | Bacteroidota | Bacteroidia | Flavobacteriales | Flavobacteriaceae | MAG-121220-bin8 | MAG-121220-bin8 sp002700465 |
| MAXBIN__023 | 307720 | 215 | 1307 | 37.5 | 23.94 | 14.08 | Bacteria | | | | | | |
| MAXBIN__015 | 102878 | 67 | 1501 | 41.1 | 23.94 | 4.23 | Bacteria | Verrucomicrobiota | Lentisphaeria | Lentisphaerales | Lentisphaeraceae | Lentisphaera | Lentisphaera araneosa |
| MAXBIN__038 | 1660543 | 1282 | 1246 | 67.7 | 22.54 | 9.86 | | | | | | | |
| MAXBIN__010 | 993817 | 673 | 1409 | 40.5 | 22.54 | 21.13 | Bacteria | Proteobacteria | Gammaproteobacteria | PS1 | Thioglobaceae | Thioglobus | Thioglobus singularis |
| MAXBIN__006 | 838719 | 546 | 1501 | 41.4 | 22.54 | 8.45 | Bacteria | | | | | | |
| MAXBIN__004 | 123978 | 69 | 1799 | 35.6 | 21.13 | 1.41 | Bacteria | Marinisomatota | Marinisomatia | Marinisomatales | TCS55 | TCS55 | TCS55 sp001577025 |
| MAXBIN__037 | 1869842 | 1462 | 1221 | 66.9 | 0.00 | 0.00 | | | | | | | |
| MAXBIN__008 | 1481191 | 950 | 1536 | 42.0 | 0.00 | 0.00 | | | | | | | |
| MAXBIN__029 | 733196 | 440 | 1500 | 35.9 | 0.00 | 0.00 | Bacteria | | | | | | |
| MAXBIN__002 | 668219 | 440 | 1498 | 46.9 | 0.00 | 0.00 | Bacteria | Proteobacteria | Gammaproteobacteria | Enterobacterales | Vibrionaceae | Aliivibrio | Aliivibrio salmonicida |
| MAXBIN__028 | 635327 | 444 | 1369 | 47.3 | 0.00 | 0.00 | Bacteria | | | | | | |
| MAXBIN__017 | 499776 | 310 | 1538 | 35.0 | 0.00 | 0.00 | Bacteria | Bacteroidota | Bacteroidia | Flavobacteriales | BACL11 | UBA8444 | UBA8444 sp003454845 |
| MAXBIN__001 | 248620 | 152 | 1585 | 46.4 | 0.00 | 0.00 | | | | | | | |
| MAXBIN__012 | 239179 | 165 | 1417 | 31.5 | 0.00 | 0.00 | | | | | | | |
| MAXBIN__005 | 207317 | 135 | 1573 | 36.4 | 0.00 | 0.00 | Bacteria | Proteobacteria | Gammaproteobacteria | PS1 | Thioglobaceae | Thioglobus | Thioglobus singularis |
| MAXBIN__027 | 155301 | 122 | 1174 | 34.0 | 0.00 | 0.00 | | | | | | | |
| MAXBIN__020 | 121882 | 81 | 1527 | 31.4 | 0.00 | 0.00 | Bacteria | Bacteroidota | Bacteroidia | Flavobacteriales | Flavobacteriaceae | MED-G11 | |
| MAXBIN__030 | 102295 | 53 | 1953 | 33.1 | 0.00 | 0.00 | Bacteria | Bacteroidota | Bacteroidia | Flavobacteriales | Flavobacteriaceae | Maribacter | Maribacter sp000153165 |
